# Target gene responses differ when transcription factor levels are acutely decreased by nuclear export versus degradation

**DOI:** 10.1101/2024.05.20.595009

**Authors:** James McGehee, Angelike Stathopoulos

**Affiliations:** California Institute of Technology, Division of Biology and Biological Engineering, 1200 East California Boulevard, Pasadena, CA 91125

## Abstract

Defining the time of action for morphogens requires tools capable of temporally controlled perturbations. To study how the transcription factor Dorsal affects patterning of the *Drosophila* embryonic dorsal-ventral axis, we used two light-inducible tags that result in either nuclear export or degradation of Dorsal when exposed to blue light. Nuclear export of Dorsal results in loss of expression for the high threshold, ventrally-expressed target gene *snail* (*sna*) but retention of the low threshold, laterally-expressed target gene *short-gastrulation* (*sog*). In contrast, degradation of Dorsal results in retention of *sna,* loss of *sog*, and lower nuclear levels than when Dorsal is exported from the nucleus. To elucidate how nuclear export results in loss of *sna* but degradation does not, we investigated Dorsal kinetics using photobleaching and found it reenters the nucleus even under conditions of blue-light when export is favored. The associated kinetics of being imported and exported continuously are likely responsible for loss of *sna* but, alternatively, can support *sog*. Collectively, our results show that this dynamic patterning process is influenced by both Dorsal concentration and nuclear retention.

**SUMMARY STATEMENT:** This study shows how optogenetic tools can be used to determine how a transcription factor’s levels and nuclear retention impact a dynamic patterning process.

## INTRODUCTION

Morphogens are proteins that form concentration gradients in developing organisms to regulate the expression of target genes (Kicheva and Briscoe 2023; Rogers and Schier 2011). The response of target gene expression to particular morphogen concentrations is termed a threshold response. Only when levels of the morphogen are over a particular concentration is expression supported. Through threshold responses, morphogen concentration controls target gene expression to pattern fields of cells, such that distinct spatial domains express different genes. This patterning process ultimately helps to define distinct tissues. Morphogens can be extracellular signaling molecules, such as sonic hedgehog (SHH) and bone morphogenic protein (BMP), as well as intracellular transcription factors that form a concentration gradient such as Bicoid and Dorsal in the syncytial *Drosophila* embryo (Briscoe and Small 2015; Greenfeld, Lin, and Mullins 2021).

While the spatial regulation of morphogen gradients has been well studied and it has long been appreciated that patterning is a dynamic process (Kutejova, Briscoe, and Kicheva 2009; Nahmad and Lander 2011; Rushlow and Shvartsman 2012), the time of action of morphogen gradients has been more difficult to investigate. Some studies of morphogens have shown that morphogen signals can be integrated over time to support expression of target genes, showing that duration of signal is important (Sagner and Briscoe 2017; Huang et al. 2017). However, it is not always possible to determine the temporal role of a morphogen using genetic knockouts and epistasis analyses. If a factor has multiple roles in a gene regulatory network, loss of the factor through genetic perturbations often prevents the gene regulatory network from reaching subsequent steps. Thus, techniques that can remove a protein with fine spatiotemporal control are needed to study a factor’s role at multiple steps in a network. Optogenetics, or the use of light to control genetically encoded proteins, allows for the study of these interactions and dynamics, and has proven useful in studying morphogen patterning (Rogers et al. 2020; Huang et al. 2017; McDaniel et al. 2019; Singh et al. 2022; Johnson et al. 2017). Here, we have used two optogenetic tags, blue light inducible degradation domain (Bonger et al. 2014) and a light inducible nuclear export system (Niopek et al. 2016), to determine if different types of perturbations to the nuclear levels of the transcription factor Dorsal (DL) differentially affect target gene expression in *Drosophila melanogaster* embryogenesis.

The early stages of *Drosophila* embryogenesis consist of rapid DNA replication and nuclear divisions, called nuclear cycles (nc). Many zygotic genes are activated during nc14, which is the longest nuclear cycle during early embryogenesis. During this time, DL forms a nuclear concentration gradient across the dorsal-ventral (DV) axis leading to expression of high threshold target genes in ventral regions and low threshold target genes more dorsally, in ventrolateral and lateral regions (Reeves and Stathopoulos 2009). While DL levels are instrumental in specifying target gene expression in a spatially-instructive manner consistent with morphogen-outputs, nuclear DL levels are also dynamic and increase during these early stages (Rushlow and Shvartsman 2012) raising the questions when and for how long does DL concentration have an effect on target gene expression.

In a previous study, we used BLID to degrade DL at certain points in time to understand how the loss of nuclear DL during certain nuclear cycles played a role in target gene expression (Irizarry et al. 2020). In that study, we found that *snail* (*sna*), which is thought to require high nuclear DL levels, can be activated independently of high DL levels in late nc14. Presumably, high DL levels are required earlier but once the DL-target gene Twist (Twi) is expressed, high DL levels are no longer required. In support of this model, we found that this late activation of *sna* requires Twi. While BLID is able to remove a transcription factor in a spatiotemporally controlled manner under blue light illumination, the system is only reversible if nascent protein is made after the light is removed and, for DL, no nascent DL-BLID is made during these stages of embryogenesis. Therefore, DL-BLID could define at what time DL stops being required for *sna* expression (i.e. at late 14), but using this system we were not able to separate DL’s early temporal role from its later role.

For this purpose, LEXY (Niopek et al. 2016; Kögler et al. 2021; Singh et al. 2022; Zhao et al. 2023), which allows for the reversible depletion of nuclear protein through blue-light inducible nuclear export was added to DL. To test the effects of depleting DL on target gene expression, nascent transcription was imaged live using the MS2/MCP system (H. G. Garcia et al. 2013; Lucas et al. 2013). Combining BLID, LEXY, and live imaging, we discovered that the *sna* regulatory system responds differently to loss of nuclear DL when comparing LEXY to BLID. Despite DL-LEXY being associated with higher nuclear DL levels than DL-BLID, *sna* is lost with LEXY but retained with BLID under blue light. This difference in responding to the loss of DL suggests that import-export kinetics influence target gene expression and are an important regulatory factor, in addition to concentration, for patterning the DV axis.

## RESULTS

### Dorsal is exported from the nucleus under blue light in *dl-LEXY*

To develop a system where we could control nuclear Dorsal levels reversibly, we used CRISPR/Cas9 genome editing to construct in-frame fusions of DL and DL-mCherry to the LEXY tag (Niopek et al. 2016), generating the *Drosophila* stocks *dl-LEXY* and *dl-mCh-LEXY* (Fig. 1A,B ‘LEXY’). These stocks are homozygous viable and fertile in the dark. When embryos expressing DL-LEXY are treated with blue light, it results in export of the fusion protein from the nucleus as detected by following the mCherry signal associated with DL-mCh-LEXY during live imaging (Fig. 1B-D; Movie 1). After the blue light is removed, DL-mCh-LEXY again enters nuclei within 1-5 minutes (Movie 1) and levels recover (Fig. 1C,D). This process of export and reimport can be repeated multiple times (Movie 1), which makes it possible to use live imaging to test how temporarily removing DL affects target gene expression. We sought to test whether two different approaches to perturb DL levels, transiently with DL-LEXY or permanently with DL-BLID, lead to different gene expression outcomes.

**Figure 1.**
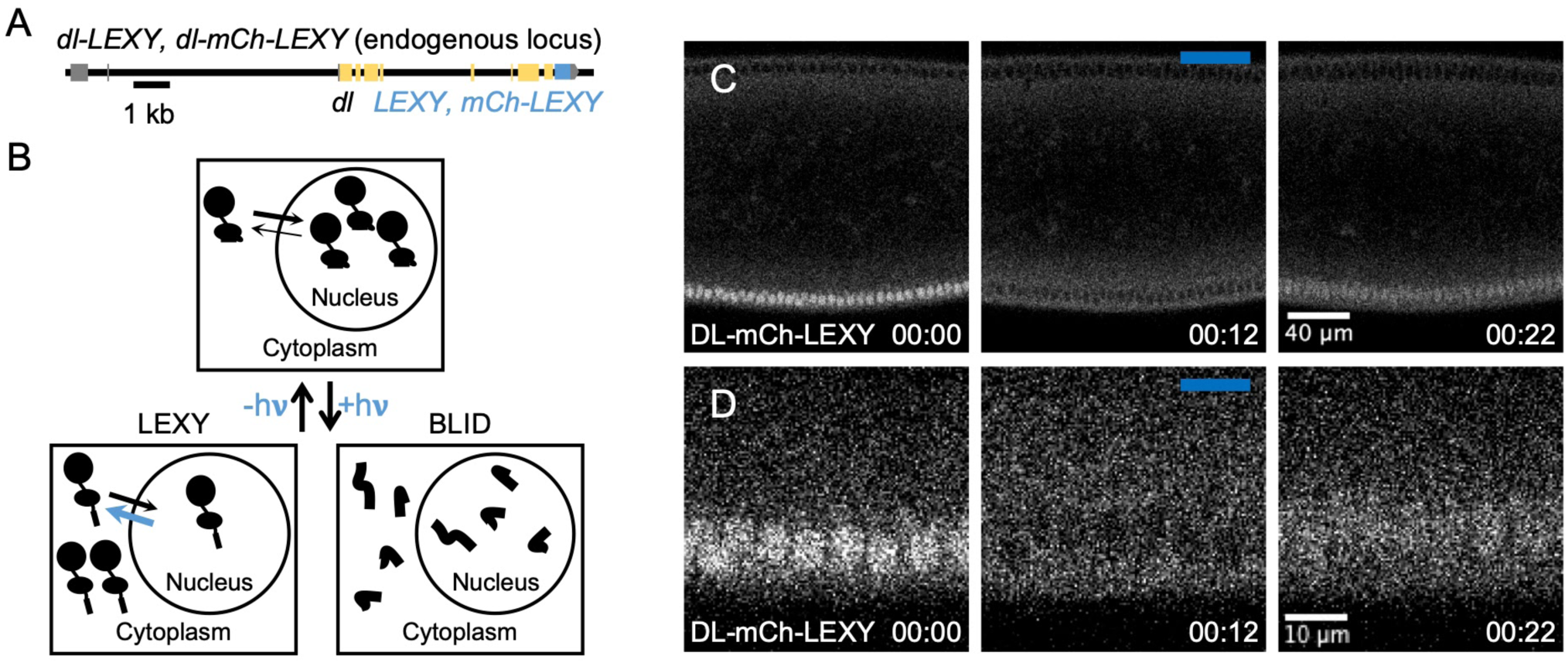
Dorsal is exported from the nucleus under blue light in *dl-LEXY*. (**A**) A schematic of the *dl-LEXY* and *dl-mCh-LEXY* CRISPR constructs. **(B)** Blue light reveals a NES in LEXY, leading to nuclear export, or a degron in BLID, leading to degradation. **(C)** Stills during nc14 from a movie of DL-mCherry-LEXY before (00:00), during (00:12), and after (00:22) blue light exposure. Time stamps represent the amount of time since image acquisition began. The blue bar represents images taken under blue light. (D) The same images as in C, only zoomed in. Embryos in C and D are oriented with the anterior (A) to the left, posterior (P) to the right, dorsal (D) at the top, and ventral (V) at the bottom. For DL-mCh-LEXY n = 2. Embryos were collected from different cages but from the same stock.

### Sites of active *sna* transcription are lost in *dL-LEXY* but recover in *dL-BLID* when DL is removed with blue light at mid-nc14

DL activates expression of an array of target genes including the high threshold target *snail* (*sna*) in the ventral region (Fig. 2A,B). To test the effect of removing DL transiently at mid-nc14, when nuclear concentration of DL in ventrally-positioned nuclei is highest (Reeves et al. 2012), embryos from *dl-LEXY* mothers were illuminated with blue light for 10 min or 20 min and compared to embryos that were kept in the dark during a similar period of time (Fig. 2C). To monitor expression of *sna*, we used a previously published MS2 reporter that was inserted at an exogenous location (Fig. S1A; Bothma et al. 2015; Irizarry et al. 2020). Using the MS2-MCP imaging system, we were able to observe expression of *sna* over time from a ventral viewpoint (Fig. 2D). We previously found that *sna* transcription at later stages of nc14 can be independent of high levels of DL, as nascent transcription is detected in embryos laid by *dl-BLID* mothers that develop under blue light at mid/late-nc14 (Irizarry et al. 2020). To compare *dl-LEXY* and *dl-BLID*, we reimaged *dl-BLID* under the same conditions as *dl-LEXY*.

**Figure 2.**
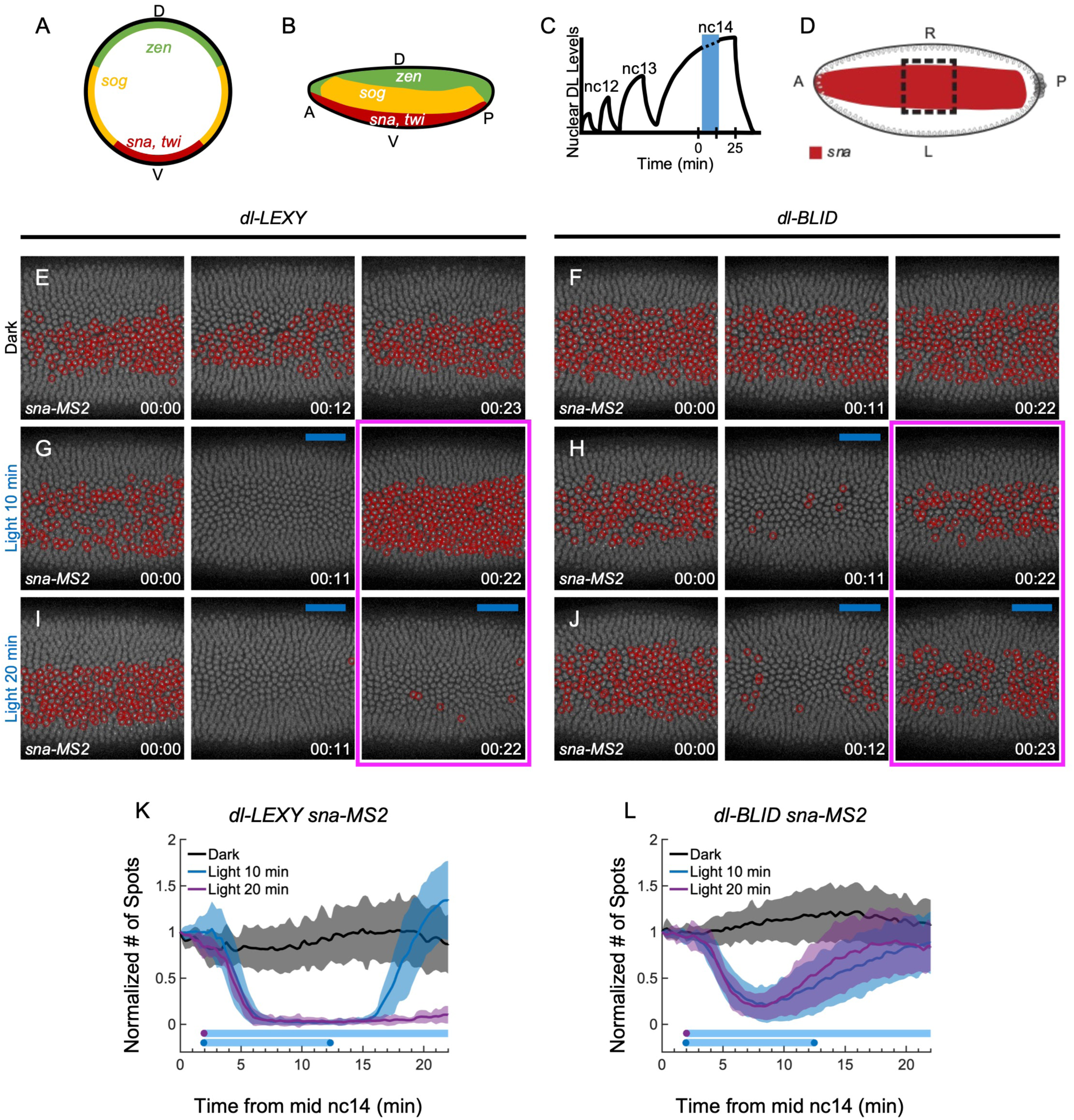
Sites of active *sna* transcription are lost in *dL-LEXY* but recover in *dL-BLID* when DL is removed with blue light at mid-nc14. (**A,B**) Expression of DL target genes along the DV axis in a cross section (A) or lateral view (**B**) with high threshold targets (*sna* and *twi*; red), low threshold targets (*sog*; yellow), and repressed targets (*zen*; green). **(C)** Illumination scheme relative to schematic of nuclear DL concentration trends over time from previously quantified data (Reeves et al. 2012). Embryos were illuminated for 10 min with blue light during mid-nc14. **(D)** A schematic of the *sna* domain (red) in a ventral view. The black dashed box represents the field of view and the area of blue light illumination. **(E, G, I)** *sna-MS2* expression in *dl-LEXY* during late nc14, when kept in the dark (E), illuminated for 10 min (G), or illuminated for 20 min (I), at 0 min (00:00), 11 min (00:11 and 00:12), and 22 min (00:22 and 00:23). **(F, H, J)** Similar conditions to E, G, and I, except for *dl-BLID*. **(K,L)** The normalized mean number of spots, or transcription foci, (mean ± s.d., n = 6 for each, except for dl-LEXY light 20 min, n = 7) detected for *sna-MS2* in *dl-LEXY* (E, G, I) and *dl-BLID* (F, H, J) over time. For each embryo, the mean number of spots was divided by the starting number of spots to normalize the data. Black is in the dark, blue is 10 min illumination, and purple is 20 min illumination. The blue bars represent the average illumination window for the matching condition. For ventral views (i.e. E-J), embryos are oriented with the anterior (A) to the left, posterior (P) to the right, and ventral (V) facing out of the page, and the field of view is positioned in the center of the trunk. Foci are circled in red for *sna*. t=0 indicates the start of imaging, at mid-nc14, as determined by the cellularization front having progressed 50% of nuclei length. In this and all other figures, blue bars represent frames under blue light and time stamps are hours:minutes. Embryos of a certain genotype were collected from the same cage either on the same day or subsequent days.

In the dark, sites of active transcription are detected continuously for *sna* in ventral nuclei during mid-nc14 in *dl-LEXY* (Fig. 2E; Movie 2) and *dl-BLID* (Fig. 2F; Movie 2). After 10 min of blue light exposure, sites of nascent *sna* transcription are undetectable in most nuclei in *dl-LEXY* (Fig. 2G 00:11; Movie 2) or are detected in only a few nuclei in *dl-BLID* (Fig. 2H 00;11; Movie 2). In both *dl-LEXY* and *dl-BLID*, sites of active *sna* transcription are detected after returning to the dark (Fig. 2G,H 00:22; Movie 2). Under 20 min of blue light exposure, sites of nascent *sna* transcription remain undetected in *dl-LEXY* (Fig. 2I 00:11 and 00:22; Movie 2). In *dl-BLID*, the number of active *sna* transcription sites initially decreases before increasing with 20 min of blue light (Fig. 2J 00:12 and 00:23; Movie 2), similar to the case of 10 min of blue light (Fig. 2H). These results agree with our previous study for *dl-BLID* (Irizarry et al. 2020), however *dl-LEXY* results in a different outcome. Specifically, these data demonstrate that nuclear DL is required to maintain sites of active *sna* transcription at mid/late-nc14 in *dl-LEXY* but not in *dl-BLID*.

At least six movies were obtained for each condition: dark, 10 min blue light, and 20 min blue light, for both *dl-LEXY* and *dl-BLID*. To quantify the changes in the number of active transcription sites, foci were detected and counted over time. Upon blue light illumination, the quantification shows the initial decrease in the number of active transcription sites in both *dl-LEXY* and *dl-BLID* (Fig. 2K,L, t=4 min), and how this number increases in *dl-BLID*, but not *dl-LEXY*, at later time points when blue light remains on (Fig. 2K,L, t=12 min and 22 min). The number of active *sna* transcription sites in *dl-BLID* increases similarly after ∼10 min, regardless of whether the embryo was exposed to blue light for 10 min or 20 min (Fig. 2L). To determine if there was any difference in the starting number of active *sna* transcription sites, we compared the mean starting number of transcription sites between *dl-BLID* and *dl-LEXY*. At the starting point, no light has been applied, so all conditions are comparable. We found no significant difference in the mean number of starting *sna* transcription sites (Fig. S2A; *dl-BLID* vs *dl-LEXY*, p = 0.5, n = 19 for *dl-LEXY* and n = 18 for *dl-BLID*), demonstrating that the starting number of transcription sites does not explain the difference we see between *dl-BLID* and *dl-LEXY*. These results suggest that blue light illumination results in different trends in *sna* transcription for *dl-BLID* versus *dl-LEXY*, showing that transcription can recover when DL is degraded (BLID) but not when it is exported (LEXY) (Fig. 2K,L).

### In *dl-LEXY*, *sog* expression is supported even under blue light

Since *dl-LEXY* and *dl-BLID* result in a different number of active *sna* transcription sites after blue light illumination, we sought to determine if the phenomenon was unique to *sna*, or if other targets of DL behaved similarly. One such target gene is *short gastrulation* (*sog*), which is a low threshold target and is expressed broadly in lateral regions (rev. in Reeves and Stathopoulos 2009). Sna acts as a transcriptional repressor to repress *sog* expression in the ventral domain, refining *sog* expression to two lateral stripes. DL also acts directly as a repressor to limit the expression of the gene *zerknüllt* (*zen*) to dorsal regions. Expression of *zen* is thought to respond to a similar threshold as *sog*, which results in *zen* having an inverse pattern compared to *sog* (Fig. 2A,B). To test the effects of changing nuclear DL levels using *dl-LEXY* and *dl-BLID* on these target genes, we created *sog-MS2* and *zen-MS2* at their endogenous loci using CRISPR/Cas9 (Fig. S1A). Previously, changes in the *sog* boundary in fixed samples were only detected if we illuminated *dl-BLID* embryos for a prolonged period (Irizarry et al. 2020). Thus, we illuminated continuously from the end of nc12 until germ-band extension was observed (i.e. all of nc13 and nc14, Fig. 3B).

**Figure 3.**
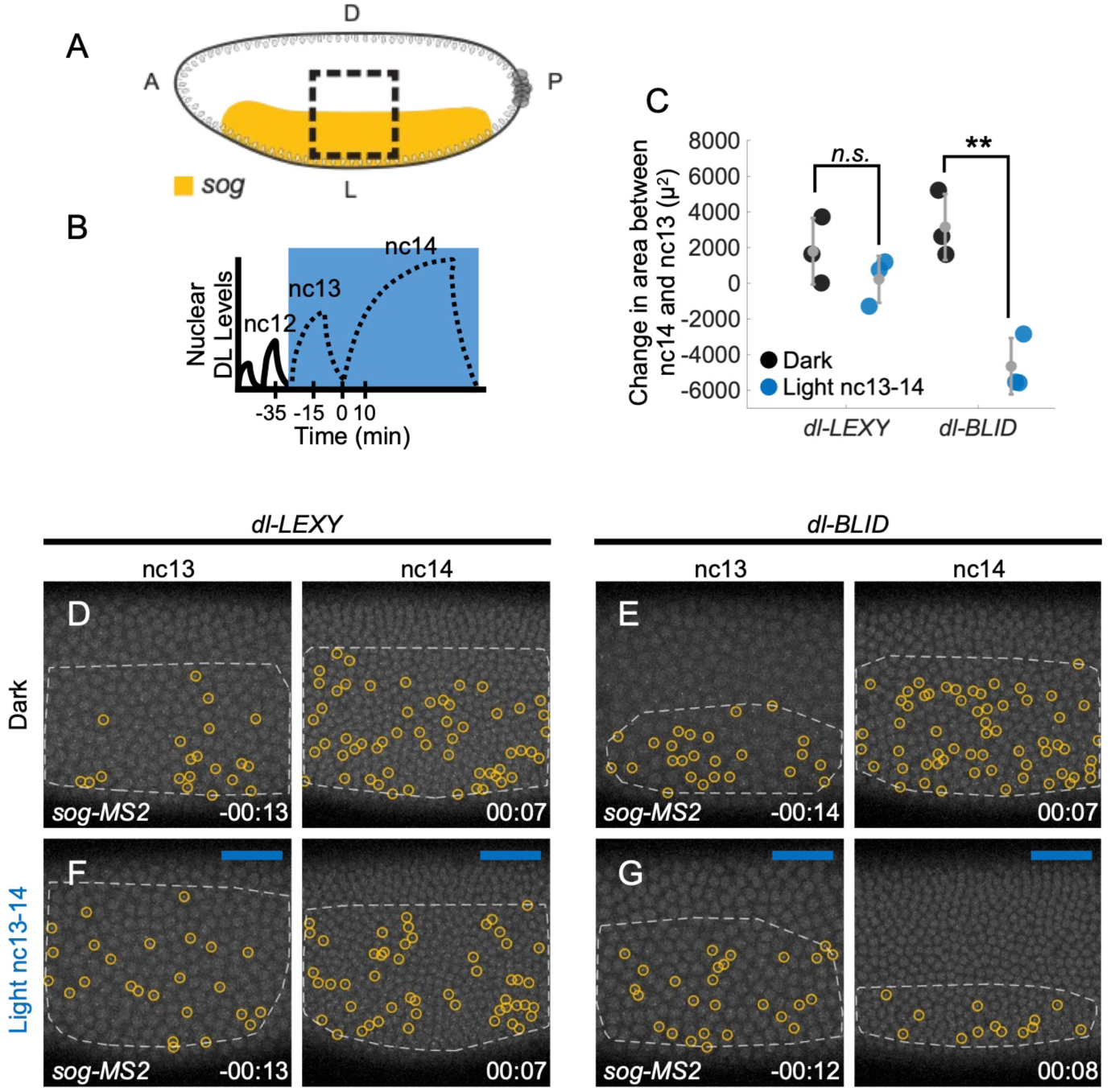
In *dl-LEXY*, *sog* expression is supported even under blue light. (**A**) A schematic of the *sog* domain (gold/yellow) transcription in a dorsal-lateral view. The dashed black box represents the field of view and the area of blue light illumination. **(B)** Illumination scheme displayed relative to nuclear DL concentration trends over time (schematic). Embryos were illuminated with continuous blue light in nc13 through nc14. **(C)** Quantification of the change in area between nc14 and 13 for *sog-MS2* in *dl-LEXY* and *dl-BLID*. Black markers represent the dark and blue markers represent illumination from nc13-14 (in gray, mean ± s.d., n = 3 for each). When comparing dark to light at nc13-14, only *dL-BLID* is significantly different, p = 0.002 (Tukey’s HSD for multiple comparisons after performing one way ANOVA). **(D, F)** *sog-MS2* in *dl-LEXY* when kept in the dark (D) or when illuminated continuously from nc13 through nc14 (F) at nc13 (–00:13) and early-nc14 (00:07). **(E,G)** Similar to D,F, except for *dl-BLID*. Foci are circled in yellow for *sog*. t=0 indicates the start of nc14. Embryos of a certain genotype were collected from the same cage either on the same day or subsequent days.

To detect differences in the *sog* and *zen* expression domains, we imaged in a dorsal-lateral field of view, which captures the dorsally-positioned boundary of *sog* and the boundary of *zen* and quantified the change in domain between nc13 and nc14 (Fig 3A,C). For *sog-MS2*, signal was detected in the dark at nc13 and nc14 in *dl-LEXY* (Fig. 3D, Movie 3) and *dl-BLID* (Fig 3E, Movie 3). When illuminated during nc13-14, there was not a significant difference in the position of the dorsal boundary of the *sog* domain in *dl-LEXY* (Fig. 3C,F; Movie 3), but there was a retraction (i.e. less spots detected in field of view) in the *sog* boundary in *dl-BLID* (Fig. 3C,G, p = 0.002; Movie 3). In contrast to *sog*, the *zen* boundary position remained unchanged by any of these similar perturbations (Fig. S3A-G; Movie 6; see Discussion).

### Under blue light activation DL levels are equal or higher in *dl-LEXY* compared to *dl-BLID*

To potentially explain the differences we observed in *dl-LEXY* and *dl-BLID* for *sna* and *sog,* we assayed the levels of nuclear DL in these lines under blue light. In the traditional threshold model, *sna* expression requires the highest levels of DL (Fig. 4A). In this model, we would predict that DL levels are higher in *dl-BLID* than in *dl-LEXY* when under blue light. To test this, embryos laid by *dl-mCh-LEXY* and *dl-mCh-BLID* mothers were illuminated with 10 min and 20 min of blue light (Fig. 4B), as done in the previous *sna* experiment (Fig. 2). In both *dl-mCh-LEXY* and *dl-mCh-BLID* in the dark, there is continuous nuclear DL (Fig. 4C,D; Movie 4). Upon 10 min of blue light illumination, DL-mCh-LEXY is exported out of the nucleus, resulting in a decrease in nuclear DL levels and an increase in cytoplasmic DL levels (Fig. 4E 00:11; Movie 4). Upon returning to the dark, nuclear DL levels begin to increase. Nuclear DL levels in the dark after being illuminated for 10 min are similar to the nuclear DL levels of embryos kept in constant darkness (Fig. 4C,E 00:22; Movie 4). Under 10 min blue light, DL-mCh-BLID levels decrease in both the cytoplasm and the nucleus. This trend continues even after the blue light is removed, suggesting that there is a slight delay in the degradation of DL (Fig. 4F; Movie 4). Upon 20 min of blue light illumination, DL-mCh-LEXY stays cytoplasmic and maintains low levels of nuclear DL (Fig. 4G; Movie 4). Under 20 min of blue light, DL-mCh-BLID levels decrease even more than the 10 min exposure, as expected (Fig. 4H; Movie 4).

**Figure 4.**
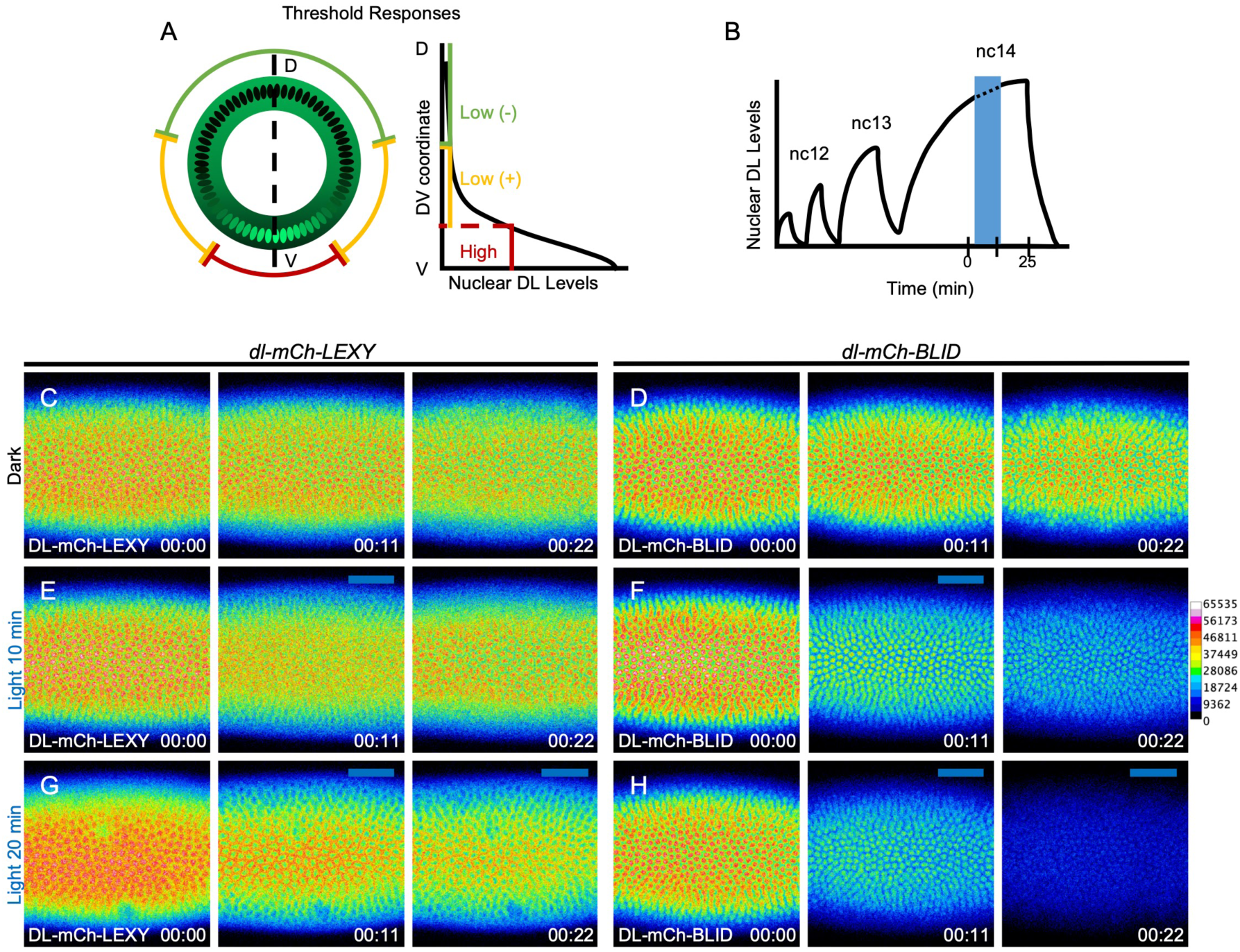
Under blue light activation, DL levels are equal or higher in *dl-LEXY* compared to *dl-BLID*. (**A**) A model of threshold responses to the DL gradient, shown in cross-section embryo view where the highest levels of nuclear DL correspond to ventral genes (red), and lower nuclear DL levels correspond to lateral genes (yellow). **(B)** Illumination scheme: embryos were illuminated with 10 or 20 min of blue light in mid-nc14. **(C, E, G)** Stills from live imaging of DL-mCh-LEXY during late nc14, when kept in the dark (C), illuminated for 10 min (E), or illuminated for 20 min (G), at 0 min (00:00), 11 min (00:11), and 22 min (00:22), n = 1 for each. **(D, F, H)** Similar conditions to C, E, and G, except for DL-mCh-BLID, n = 3 for each. Images are displayed using a rainbow colormap, where high intensity pixels are in white and pink, and low intensity pixels are in blue and black. Embryos are oriented with the anterior (A) to the left, posterior (P) to the right, and ventral (V) facing out of the page, and the field of view is positioned in the center of the trunk. t=0 indicates the start of imaging, at mid-nc14, as determined by the cellularization front having progressed 50% of nuclei length. Embryos of a certain genotype were collected from the same cage either on the same day or subsequent days.

These data show that the levels of nuclear DL in DL-mCh-BLID appear lower than the levels in DL-mCh-LEXY when both are under 20 min of blue light. In contrast, the levels of nuclear DL under 10 min of blue light are similar in *dl-LEXY* and *dl-BLID*. While the difference in nuclear DL levels between *dl-LEXY* and *dl-BLID* can explain the differences we see in *sog* expression, it fails to explain the difference in the number of *sna* transcription sites between *dl-LEXY* and *dl-BLID* observed under 20 min of blue light. Since DL is a positive input to *sna*, it would be expected that *sna* expression would occur where DL levels are higher, which would be in *dl-LEXY*. However, we find that there are a greater number of *sna* transcription sites in *dl-BLID* where DL levels are lower.

### Twi levels do not change at late nc14 under blue light in *dl-LEXY* or *dl-BLID*

Since nuclear DL levels cannot explain the differences we see between *dl-LEXY* and *dl-BLID*, another possible explanation is that blue light induced export of DL may have an effect on Twi protein levels. We have previously shown that Twi was responsible for DL-independent activation of *sna* at late nc14 in *dl-BLID* under blue light (Irizarry et al. 2020). To test whether Twi levels were differentially affected in *dl-LEXY* and *dl-BLID* under blue light, we imaged Twi using a previously published Twi-LlamaTag fly stock (Bothma et al. 2018), which allows us to detect Twi protein localization live in vivo when maternally deposited mCherry is available. Using Twi-LamaTag, we found that the levels of Twi do not change upon 20 min blue light illumination and appear similar between *dl-LEXY* and *dl-BLID* (Fig. S4). This suggests that a difference in Twi levels cannot explain the difference in *sna* expression.

An additional possible explanation for how DL-LEXY and DL-BLID differentially affect *sna* expression is that DL-LEXY sequesters potential cofactors in the cytoplasm under blue light. Implicitly, this means that when DL is exported from the nucleus, any transcription factors or cofactors bound to DL would also shuttle into the cytoplasm. There is some evidence that DL and Twi might physically interact (Shirokawa and Courey 1997). If Twi was exported with DL under blue light, we should see a change in localization of the Twi protein using the Twi-LlamaTag; however, we did not observe export of Twi protein under blue light in *dl-LEXY* (Fig. S4C; Movie 7). This lack of reduction in Twi nuclear levels upon blue light illumination is especially clear when comparing the nuclear levels of Twi to the blue light induced export of DL-mCh-LEXY (Fig. S4C compared to Figs. 1C,D and 4E,G; Movie 7 compared to Movie 1 and Movie 4). This demonstrates that DL nuclear export does not result in an appreciable reduction of nuclear Twi.

While Twi is not exported with DL, this does not rule out that other potential cofactors might be exported with DL in *dl-LEXY* under blue light. For instance, since Twi is required for *sna* transcription in *dl-BLID* under blue light, it is possible that DL and Twi share a cofactor, and it is this cofactor that is sequestered with DL in the cytoplasm in *dl-LEXY* under blue light. However, it would be difficult to test the protein localization of every DL cofactor individually, especially if the cofactor in question is unknown. Thus we sought to explore and rule out other possible explanations for how *dl-LEXY* and *dl-BLID* result in differences in the number of *sna* transcription sites.

### FRAP demonstrates DL-LEXY transits in and out of the nucleus, even under blue light illumination

Another model that could explain the differences between *dl-LEXY* and *dl-BLID* relates to the dynamics of how nuclear protein levels decrease with these perturbations. In *dl-LEXY* under blue light, DL is likely constantly transiting in and out of the nucleus, while in *dl-BLID*, DL is either degraded or remains nuclear. When DL-LEXY is exposed to blue light, the normally buried nuclear export sequence (NES) is revealed as the J⍺ helix is unfolding (Niopek et al. 2016). However, DL’s nuclear localization sequence (NLS) should not be affected. Thus, DL would be constantly transiting between the cytoplasm and the nucleus, being biased to the cytoplasm due to the strong nature of the exposed NES contained in LEXY. We hypothesized that this constant import and export may act to disrupt DL-independent activation of *sna*. Specifically, DL might act transiently at the *sna* locus before being exported, causing a disruption in *sna* activation through other factors, such as Twi. This model suggests that *sna* needs DL to remain in the nucleus for a sustained amount of time.

To test if DL is transiting in and out of the nucleus, we performed fluorescence recovery after photobleaching (FRAP) on DL-mCh-LEXY under blue light. Using FRAP, we bleached a large region of interest (ROI), shown by the green circle, and quantified the nuclear levels in the center most bleached nucleus using the yellow ROI (Fig. 5A). The quantification reveals that nuclear DL-mCh-LEXY signal recovers after bleaching (Fig. 5B-D; Movie 5). The FRAP was done under blue light so that nuclear export is favored. FRAP of DL-mCh-LEXY under blue light demonstrates that DL-mCh-LEXY levels recover in the nucleus even when export is favored, suggesting that DL-mCh-LEXY is moving in and out of the nucleus.

**Figure 5.**
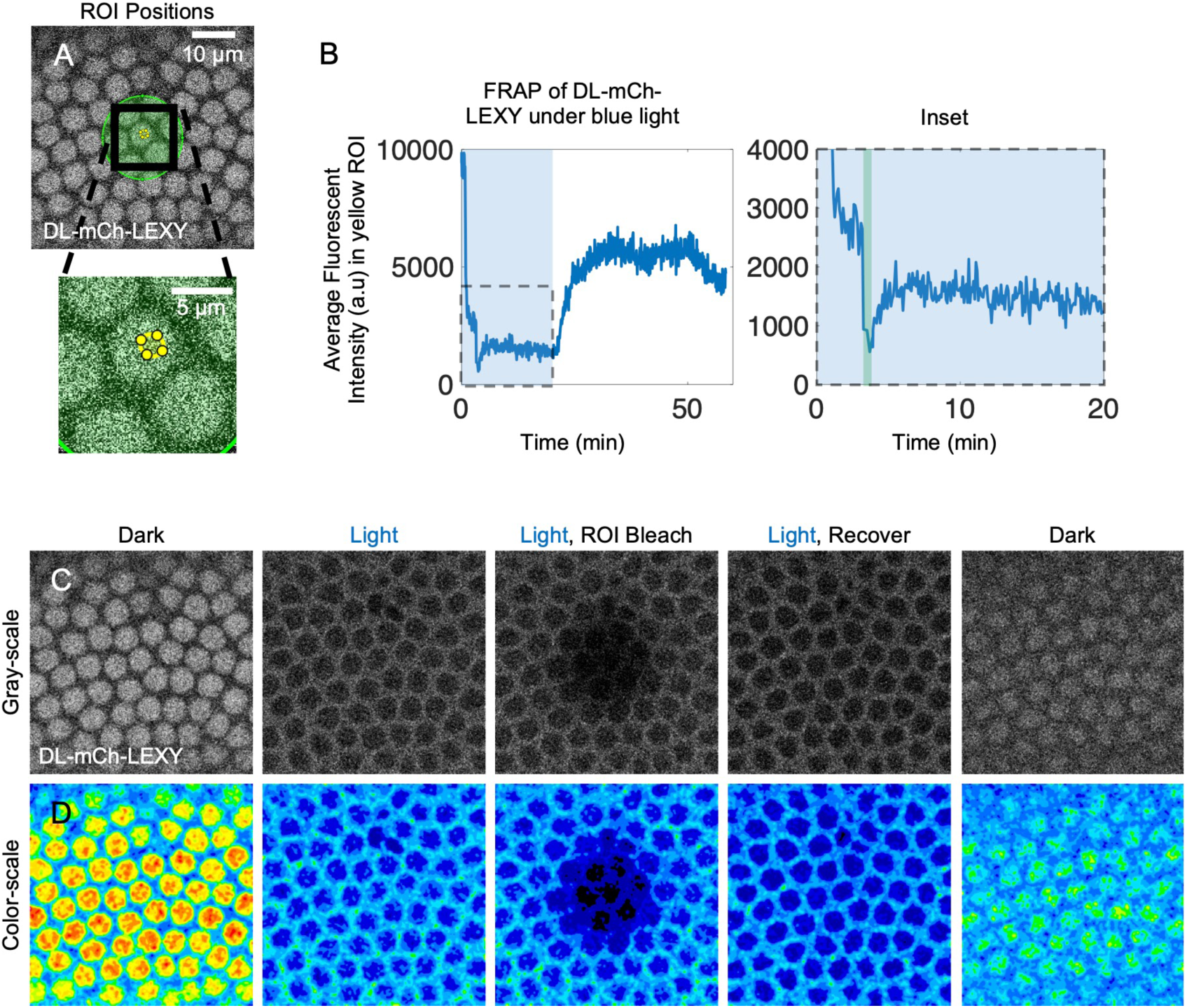
FRAP demonstrates DL-LEXY transits in and out of the nucleus, even under blue light illumination. (**A**) Location of ROIs used for bleaching (green ROI) or quantification (yellow ROI, shown in zoomed in view) during FRAP of DL-mCh-LEXY under blue light. **(B)** Quantification of fluorescence intensity in the yellow ROI in A. The shaded blue region represents the time blue light is applied. The inset from the dashed box shows the bleaching (green bar on plot) and recovery. **(C)** Stills of the embryo that underwent FRAP in the dark, after blue light was applied, after bleaching, after recovery, and after returning the embryo to the dark. **(D)** The same as in panel C except a Gaussian blur (r = 3) was performed, and the image was false colored using a rainbow colormap. Ventral views of embryos are shown. t=0 indicates the start of imaging, at mid-nc14, as determined by the cellularization front having progressed 50% of nuclei length. This experiment was repeated three times using embryos collected from different DL-mCh-LEXY CRISPR/Cas9 lines.

### Mutation of potential phosphorylation sites in DL’s C-terminal NES diminishes *sna* but has little effect on *sog*

To provide support for the idea that *sog* expression does not change in *dl-LEXY* because export cannot reduce DL levels low enough, we sought alternate ways to increase nuclear export. One way of altering the nuclear export rate is by making mutations in DL’s native C-terminal NES (NES4; Xylourgidis et al. 2006). In NES4, a single serine residue (S665) has been identified as a site of phosphorylation through a large mass spectrometry screen (Hilger et al. 2009). Since export sequences are largely hydrophobic (Kosugi et al. 2014), phosphorylation of NES4 might directly act to decrease the export rate. Other ways phosphorylation could affect the export rate, indirectly, is by masking a NES through regulation of a conformational change or interaction with a binding partner to support nuclear retention. In either case, by blocking phosphorylation, we hypothesized that cytoplasmic DL would increase due to an increase in export (Nardozzi, Lott, and Cingolani 2010). To be sure we removed all possible sites of phosphorylation in NES4, we mutated S665 and three other local serine residues. Specifically, we mutated the four serine residues to alanine residues (*dl NES S>A*) that cannot be phosphorylated or to aspartic acid residues (*dl NES S>D*) to mimic a constitutively phosphorylated state due to their negative charge (Fig. 6A). These mutations were made using large rescue constructs that include the known regulatory sequences of DL and were assayed at one copy in a *dl^1^/dl^4^* mutant background. Venus was included at the C-terminal end of these DL transgenes to support live imaging.

**Figure 6.**
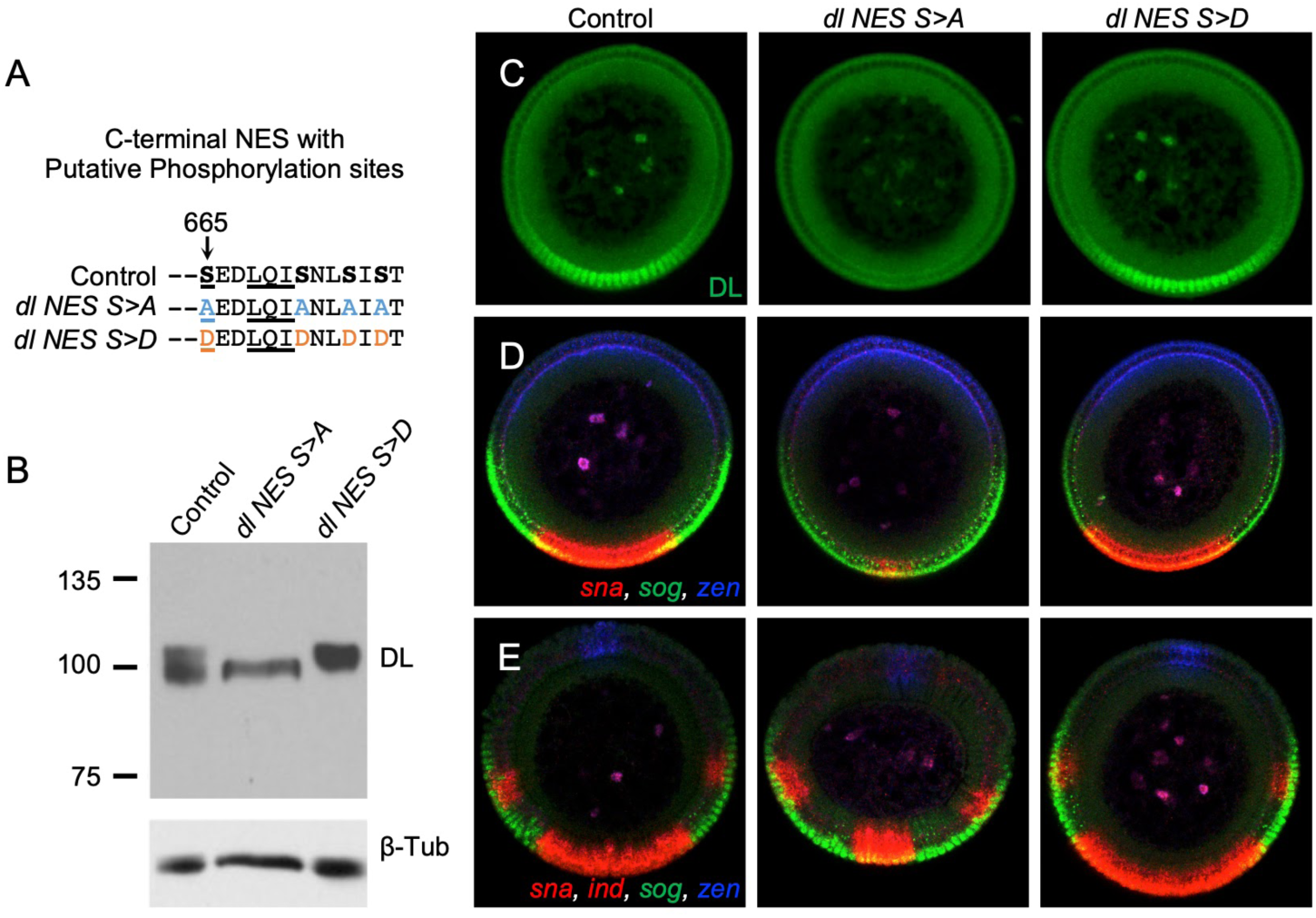
Mutation of potential phosphorylation sites in DL’s C-terminal NES diminishes *sna* but has little effect on *sog*. (**A**) Four serine residues (bold) were mutated to either four alanine (blue) or four aspartic acid (orange) residues. S665 has been shown to be phosphorylated (Hilger et al. 2009) and the underlined LQI residues have been shown to be important for the NES to function as mutation of the L and I residues result in a loss of export (Xylourgidis et al. 2006). **(B)** Western blot for DL in a *dl* null background each containing *dl-Venus* large rescue constructs: Control (*dl-Venus*)*, dl NES S>A*, and *dl NES S>D*. **(C)** Anti-DL staining in Control*, dl NES4 S>A*, and *dl NES4 S>D*. **(D,E)** In situ hybridization for *sna* and *ind* (red, same channel), *sog* (green), and *zen* (blue) in Control*, dl NES S>A*, and *dl NES S>D* at early nc14 (D) or late nc14 (E). For the DL antibody staining, the Control had n = 42, *dl NES S>A* had n = 31, and *dl NES S>D* had n = 44. For the in situ, the Control had n = 10 early and n = 2 late, *dl NES S>A* had n = 4 early and n = 2 late, and *dl NES S>D* had n = 7 early and n = 6 late.

We assayed the *dl NES S>A* and *dl NES S>D* lines using antibody staining and fluorescent in situ hybridization. Nuclear DL levels in *dl NES S>A* are clearly decreased compared to the control *dl-Venus* as shown by quantification of DL antibody staining (Figs. 6C and S5, p = 8.6 * 10^-20^, Tukey’s HSD for multiple comparisons after performing one way ANOVA). To confirm that total levels of DL were not affected, we detected DL levels in embryo extracts by western blot using DL antibody (Fig. 6B). We found that the total levels of DL are similar, although the distribution of the migrating band changed in *dl NES S>A*. This loss of differentially migrating bands supports the idea that phosphorylation is disrupted in *dl NES S>A*. The distribution of molecular weights in *dl NES S>D* was observed to shift slightly higher, although it is not as clear as the change in the *dl NES S>A* band.

To determine what effect the changes in the DL gradient had on target genes, we also performed fluorescent in situ hybridization against *sna*, *sog*, *zen*, and the laterally expressed gene *intermediate neuroblasts defective* (*ind*), at early nc14 (Fig. 6D) and late nc14 (Fig. 6E). We observed that the *sna* expression domain is clearly reduced in *dl NES S>A* at both stages (Fig. 6D,E *dl NES S>A*). *sog, zen,* and *ind* expression appear normal in the *dl NES S>A* mutant when compared to the control at either stage (Fig. 6D,E *dl NES S>A*). During early nc14, *zen* is expressed broadly, before refining until it is a narrow stripe at late nc14 (Jaźwińska, Rushlow, and Roth 1999) and expression of *ind* is only detected in late nc14, and not during early nc14 (M. Garcia et al. 2013). We observed that *zen* and *ind* undergo normal dynamics in the control and mutants. These data suggest that affecting the function of a native nuclear export sequence, by mutating putative phosphorylation sites to non-charged residues (Fig. 6A), results in decreased nuclear DL (Fig. 6C and S5 *dl NES S>A,*) that in turn is associated with a reduction in the *sna* domain (Fig. 6D,E *dl NES S>A*).

## DISCUSSION

We found that a threshold model is insufficient to explain the observed differences in *dl-LEXY* and *dl-BLID* and that, in addition to levels, the kinetics of import-export play a role in determining the expression of target genes (Fig 7A). We observed that the lowest levels of DL achievable in *dl-LEXY* under blue light are high enough for *sog* expression as expression remains on in *dl-LEXY* under blue light. This finding is also supported by the C-terminal NES serine residue mutations, which result in low nuclear DL levels but do not affect *sog*. In addition, DL levels are higher in DL-mCh-LEXY than in DL-mCh-BLID, and the *sog* boundary changes position only in *dl-BLID* under blue light. This shows that low levels of DL are instructional for *sog* expression since in *dl-BLID,* DL is degraded and likely goes below the threshold necessary for *sog* expression. Although the *sog* data supports a traditional threshold model, we found that LEXY and BLID resulted in different outcomes for *sna* expression that contradicted a threshold model. We observed that in *dl-LEXY*, *sna* expression is lost at late nc14, while in *dl-BLID*, *sna* expression can remain on (Fig. 2E-J) even though DL levels are higher in *dl-LEXY* than *dl-BLID* (Fig. 4G,H; Movie 4). These observed differences in target gene responses between *dl-LEXY* and *dl-BLID* have provided additional insights into how DL action, specifically, DL nuclear level, is interpreted by cis-regulatory systems to pattern the DV axis.

**Figure 7.**
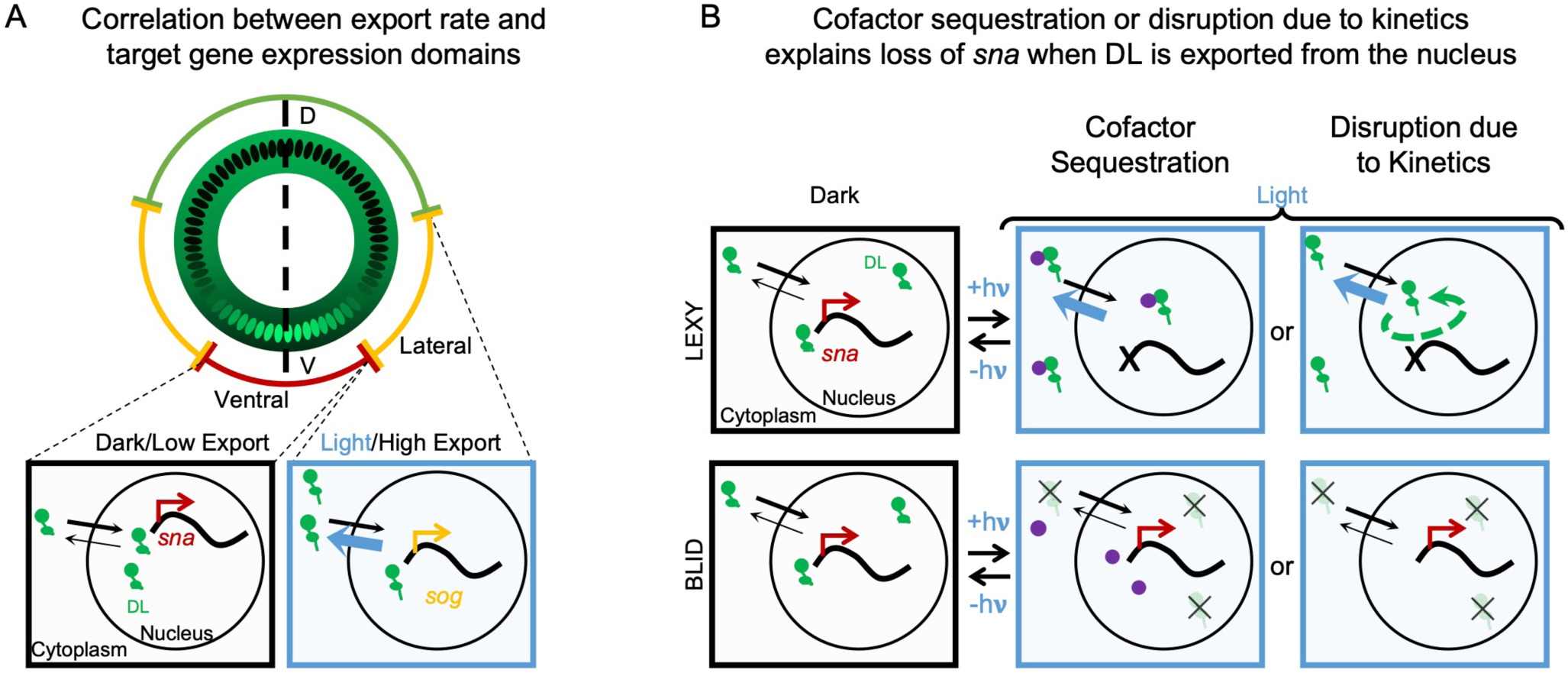
Models capable of explaining different phenotypes associated with optogenetic perturbation of nuclear DL levels through nuclear export or degradation. **(A)** Schematic of expression domains of genes (outer circle: *sna* – red, *sog* – yellow, *zen* – green) shown around an embryo cross-section displaying the DL nucleo-cytoplasmic gradient along the DV axis. Magnified views of individual nuclei from ventral (left) or lateral (right) regions are shown below and represent a working model of how low levels of export lead to sustained levels of DL sufficient for activation of *sna* (left), while high levels of export lead to low or transient levels of DL only able to activate *sog* (right). *sog* is not expressed stably in ventral regions because Sna represses it. **(B)** Two possible models to explain differences in *sna* expression detected using DL-LEXY versus DL-BLID. In the dark (first column) DL-LEXY and DL-BLID (both in green) behave similarly. In one model under blue light, DL-LEXY sequesters a cofactor (purple) in the cytoplasm, while DL-BLID does not affect the cofactor (second column). In the second model under blue light, the transient nature of DL-LEXY (green dashed arrow) blocks DL-independent activation of *sna* (red) while degradation of DL-BLID does not (third column). Thick blue arrows denote increased export rates of DL-LEXY under blue light. A thick black “**x**” marks when transcription is not activated. A thin black “x” indicates DL-BLID molecules that have been degraded by blue light.

We propose that the kinetics of DL transiting between the nucleus and cytoplasm in *dl-LEXY* disrupts *sna* expression and also instructs the patterning process (Fig. 7B “Disruption due to Kinetics”). This model is supported by the FRAP experiment, which shows that DL is transiently entering the nucleus in *dl-LEXY* under blue light (Figure 5). Since the endogenous NLS associated with DL in *dl-LEXY* is unperturbed, it follows logically that, under blue light, DL would enter the nucleus only to be exported by the strong NES in LEXY. To rule out other explanations that could explain the observed differences in *dl-LEXY* and *dl-BLID*, we compared the levels of nuclear DL in both lines under blue light. We would expect that higher nuclear DL levels in *dl-BLID* support *sna* expression, while lower nuclear DL levels in *dl-LEXY* result in a loss of *sna* expression. However, we found the opposite as DL levels are lower in *dl-mCh-BLID* than they are in *dl-mCh-LEXY* (Fig. 4); therefore, a strict concentration-dependent model does not explain the results. In addition, we evaluated whether Twi protein is exported with DL in *dl-LEXY* under blue light (Figs. S4 and 7B, “Cofactor Sequestration”). We observed that Twi is not exported with DL (Fig. S4). While it is possible that another cofactor is sequestered with DL in *dl-LEXY* under blue light, this seems unlikely because there is no evidence that DL sequesters its known interaction partner Twi. Finding neither of these alternative models sufficient to explain the observed differences, our data favors a model in which the kinetics of import-export disrupt *sna* transcription (Fig. 7B “Disruption due to Kinetics”).

While we found that *sog* responds to blue light in *dl-BLID* but not *dl-LEXY*, *zen* does not respond to DL levels in either *dl-BLID* or *dl-LEXY* when illuminated with blue light from nc13 to nc14 (Fig. S3). This is surprising because *sog* and *zen* are thought to share the same threshold, as *zen* forms an inverse pattern to *sog*. One possible explanation for why *zen* does not change is that *zen* responds to DL at a time point before we imaged. For example, *zen* may exhibit DL-dependence only between nc12 and nc13, not nc13 and nc14 when we assayed its expression domain. In addition, this might suggest that DL-repressor activity is fundamentally different from DL-activator activity, even if they are mediating similar concentration-threshold outputs. For instance, if DL binds more tightly to sites where it acts as a repressor, DL-LEXY and DL-BLID might be unable to support export or degradation, especially if degradation is primarily happening in the cytoplasm. In a future study, it would be of interest to focus on how export and degradation affect DL’s repressor activity, which may require earlier perturbations.

This study has provided general insights into the time of action of morphogens and suggests that the regulation of nuclear import-export kinetics might be important in transcription factor dynamics, providing support for recently generated models (Barros et al. 2021; Schloop, Bandodkar, and Reeves 2020). These data also suggest a general principle of how gene expression might respond to the dynamics of morphogens in addition to sensing their concentration. The transient levels needed for low threshold targets may also act to prevent transcription of high threshold targets, to reinforce the correct positioning of gene expression domains. Previous studies in yeast have focused on the encoding/decoding of information by transcription factor dynamics. With one optogenetic study of transcription factor action showing, similarly, that a subset of targets appears to be regulated by high thresholds (low affinity sites) to support high robustness (Sweeney and McClean 2023). The enhancers for *sna* are thought to contain lower affinity binding sites for DL, in comparison to enhancers for *sog* (e.g. Papatsenko and Levine 2005). Combining an analysis of binding site affinity and the pervasive use of combinatorial, cooperative binding of transcription factors using optogenetic perturbations, could be useful in understanding gene expression (Kim et al. 2024; Rao, Ahmad, and Ramachandran 2021). The expectation would be that the combinatorial control would allow genes to be less sensitive to concentration change. In support of this view, *sna* does not exhibit many linked DL and Twi sites, in contrast to DL target genes expressed in lateral regions (Markstein et al. 2004; Reeves and Stathopoulos 2009). Furthermore, studies in the field are also investigating whether stochastic gene expression is instructive or if systems are buffered against stochastic expression (Exelby et al. 2021). *sog* expression exhibits domains where expression appears stochastic (Reeves et al. 2012; MacNamara 2014) and could be a useful model for investigating how stochasticity affects gene expression.

We would argue that in addition to categorizing gene expression responses supported by morphogens in *Drosophila* and other higher animals by their threshold (high, medium, low), these responses should be categorized by their dynamical response, for example, requiring sustained, transient, or absent/reduced morphogen-dependent input. It is possible that cis-regulatory systems controlling expression of these genes might be tuned to detect sustained levels of a morphogen for high threshold targets, transient levels for low threshold targets, and absent (reduced) levels for targets that are repressed by the morphogen (rev. in Irizarry and Stathopoulos 2021). Furthermore, enhancers with different dynamical requirements have been shown to act coordinately to promote gene expression in *Tribolium* development (Mau et al. 2023). Future studies could investigate whether dynamics are sensed by enhancers and/or promoters (Hoppe et al. 2020; Delás et al. 2023). We also envision that there are other dynamic regulatory paradigms that have not yet been discovered but that optogenetic tools that provide fine spatiotemporal control will be able to help identify these paradigms and provide additional insights into how they work.

## MATERIALS & METHODS

### Fly stocks and husbandry

All *D. melanogaster* stocks were kept at 22℃ in standard medium. Experimental crosses were kept in cages with apple juice agar plates supplemented with yeast paste and were kept at 18℃. Embryos were collected for less than two weeks, such that parents were one to fourteen days old at time of collection. *w; dl-LEXY/CyO; PrDr/TM3* and *w; dl-BLID/CyO; PrDr/TM3* were crossed to *Sp/Cyo; MCP-mCherry (w+, NLS)/TM3(Bothma et al. 2018)* to generate *w; dl-LEXY/CyO; MCP-mCherry (w+, NLS)/TM3* and *w; dl-BLID/CyO; MCP-mCherry (w+, NLS)/TM3*, which were grown in bottles, and virgin *dl-LEXY; MCP-mCherry (w+, NLS)/TM3* or *dl-BLID; MCP-mCherry (w+, NLS)/TM3* were selected. These virgins were crossed to males bearing the MS2. MS2 lines included *sna-MS2 BAC (III)*, *sog-MS2 (I); Sp/Cyo, and Sp/Cyo; zen-MS2/TM3 (III)*. *sna-MS2* is a large reporter construct of ∼25kB with MS2 inserted at the 5’end of the transcript following the 5’UTR and the coding sequence replaced by the gene *yellow* (Perry et al. 2010; Bothma et al. 2015) Plasmid DNA from (Bothma et al. 2015) was inserted on the third chromosome at 65B2; 3L:6442676 (Irizarry et al. 2020). *sog-MS2* and *zen-MS2* contain insertions of MS2 within introns. In addition, *dl-mCherry-LEXY/CyO* was grown in bottles and added to experimental cages*. y^2^ cho^2^ v^1^ P{nos-phiC31\int.NLS}X; attP2 (III)* (NIG-FLY TBX-0003) was used to make *y^2^ cho^2^ v^1^ P{nos-phiC31\int.NLS}X; P{dl-gRNA}attP2 (III). y^2^ cho^2^ v^1^; Sp/CyO, P{nos-Cas9, y+, v+}2A* (NIG-FLY CAS-0004) virgins were crossed to *y^2^ cho^2^ v^1^ P{nos-phiC31\int.NLS}X; P{dl-gRNA}attP2 (III)* for injection. See Table S1 for a complete list of lines used.

### Homologous Repair Template Cloning

*LEXY* (Bonger et al. 2014) was codon optimized and, along with *MS2* (Yamada et al. 2019), synthesized by GenScript in pUC57. The *dl-LEXY*, *dl-mCh-LEXY*, *sog-MS2* and *zen-MS2* homologous repair templates were generated by editing *pHD-DsRed* (Gratz et al. 2014). The right homology arm for *dl-LEXY and dl-mCh-LEXY* was generated by PCR using a *dl-Venus-BAC(Reeves et al. 2012)* as a template, and was inserted into *pHD-DsRed* downstream of the *DsRed* using BglII and XhoI sites. The left homology arm was generated by overlap PCR, combining three fragments, the C-term of *dl*, the LEXY domain, and the *dl* 3’UTR. The left homology arm of *dl-mCh-LEXY* was made by overlap PCR, combining PCR products that used *dl-mCherry* HDR and the *dl-LEXY* HDR as a template. This PCR product was inserted into *pHD-DsRed* upstream of the *DsRed* using EcoRI and NheI sites. The *sog-MS2* homologous repair template was made by PCR, using a BAC as the template (BacPac Resource Center BACR25D05). Overlap PCR was used to mutate the gRNA binding site in the repair template. The left homology PCR product was cut with NheI and AseI and the *pHD-DsRed* plasmid was cut with NheI and NdeI to make compatible sticky ends, which were ligated together. The right homology arm PCR product and the pHD-DsRed were digested with AscI and XhoI and ligated. The *zen-MS2* homologous repair template was made the same way as the *sog-MS2* template, but used NheI and NdeI on both the insert and the backbone, and the right homology arm also used overlap PCR to mutate the gRNA sequence. The *MS2* sequence was added using NotI and AvrII, which were added to the reverse primer used to generate the left homology arm of both *sog-MS2* and *zen-MS2*. The *zen-MS2* gRNA was made by BbsI digestion of pCFD5 and Gibson assembly was used to combine the vectorized backbone and the PCR product. In both *sog-MS2* and *zen-MS2*, the MS2 sequence was inserted into the first intron, as annotated on Flybase. See Table S1 for a complete list of primers used.

### CRISPR/Cas9 Genome Editing

For *dl-LEXY* and *dl-mCh-LEXY*, *y^2^ cho^2^ v^1^; Sp/CyO, P{nos-Cas9, y+, v+}2A* virgins were crossed to *y^2^ cho^2^ v^1^ P{nos-phiC31\int.NLS}X; P{dl-gRNA}attP2 (III)*. The HDR template for *dl-LEXY* and *dl-mCh-LEXY* were injected into embryos from this cross. The *sog-MS2* HDR was co-injected with a previously made gRNA (Dunipace, Ákos, and Stathopoulos 2019) into w[1118]; PBac{y[+mDint2]=vas-Cas9}VK00027 (Bloominton #51324). For *zen-MS2*, gRNAs were found using flyCRISPR Target Finder(Gratz et al. 2014). The *zen-MS2* HDR was co-injected into *y2 cho2 v1; attP40{nos-Cas9}/CyO (NIG-FLY CAS-0001)*. For both *sog-MS2* and *zen-MS2*, Rainbow Transgenics performed the injections. All HDR templates included DsRed as a selectable marker, and transgenics were screened for DsRed expression.

### Live Imaging

Embryos from crosses between *dl-LEXY* and the *MS2* lines were collected for four hours or overnight, both at 18℃. To prepare the embryos for live imaging, embryos were hand dechorionated in the dark, using a red film (Neewer, 10087407). Embryos were transferred to an agar square and oriented so that the face that would be imaged was facing the agar. Preprepared slides were made by adding heptane glue (heptane plus double sided tape) to a coverslip that was taped to the slide and allowing it to sit overnight. This slide was used to pick the embryos up from the agar. Embryos were then checked to make sure the orientation had not been disrupted and oriented again if necessary. Water was then added to prevent desiccation of the embryos. Embryos were transferred to the microscope in a covered box. Imaging occurred on a Zeiss LMS 800 using a 25x immersible objective (LCI Plan-Neofluar 25x/0.8 Imm Korr DIC M27) at 1.7 zoom. The MCP-mCherry signal was detected using a 561 nm laser at 1% laser power with 800 V gain on a GaAsP PMT detector. The 488 nm laser at 4.5% laser power was used to perform blue light illumination with 500 V gain on a GaAsP PMT detector to protect the detector from the high laser power. Z-stacks were taken, with 30 z-planes per timepoint at 1 um thickness. Images were taken every ∼25 seconds, starting as soon as the previous z-stack finished. Images were captured as 16 bit images, and each z-slice was 512 by 512 pixels, with each pixel being 0.29 um in length and width. DL-mCh-LEXY and DL-mCh-BLID were imaged using the same settings as the MS2/MCP imaging, except the laser power of the 561 nm laser was 2% instead of 1%. Upon beginning imaging, embryos were staged under red light (white light covered with a red filter). Embryos that were the right stage and orientation were imaged. Orientation was determined by the signal being imaged. For DL-mCh-LEXY, DL-mCh-BLID, and *sna-MS2*, this meant that the signal domain was centered. For *sog-MS2* and *zen-MS2* this meant detecting the boundary of the expression domain.

### Fluorescence recovery after photobleaching (FRAP)

FRAP was imaged similarly to previous imaging setups. Images were 512 by 512 pixels, 16 bit, and taken on a Zeiss LMS 800 using a 25x immersible objective (LCI Plan-Neofluar 25x/0.8 Imm Korr DIC M27) at 5.0 zoom with a pixel size of 0.100 um. The DL-mCh-LEXY signal was detected using a 561 nm laser at 1% laser power with 850 V gain on a GaAsP PMT detector. The 488 nm laser at 4.5% laser power was used to perform blue light illumination with 500 V gain on a GaAsP PMT detector to protect the detector from the high laser power. Time points were acquired at a rate of one frame per five seconds. Bleaching was performed after 40 time points in an ROI using the 561 nm laser at 20% laser power and performing 50 iterations of bleaching at each time point. Bleaching occurred until signal intensity was 20% of original signal which occurred within three time points. The imaging scheme for FRAP to test that DL is still entering the nucleus even under blue light is as follows: we first illuminated an embryo laid by a *dl-mCh-LEXY* mother until export was observed. Then we bleached the signal in an ROI while maintaining the blue light. Once the region was bleached, we allowed the signal to recover, while still maintaining the blue light. Finally, we returned the embryo to the dark and observed an increase in nuclear DL-mCh-LEXY. To quantify the recovery inside the nucleus, we drew a small ROI inside a single nucleus, which was visually confirmed to be in the nucleus at every time point, even as the nucleus shifted positions slightly.

### Fixed Imaging

For fixed sample preparation of the *dl-Venus*, *dl NES S>A*, and *dl NES S>D*, embryos were collected for 1 hour and aged 2-3 hours and then were dechorionated in bleach, fixed in 4 mL of 9.25% formaldehyde and 4 mL of heptane for 20 minutes and then rinsed and stored in methanol at –20°C. For in situ hybridization, protocols were followed as described previously (Kosman et al. 2004) using riboprobes generated for *sna*, *ind*, *sog*, and *zen*. Sheep anti-digoxigenin (Life Technology PA185378), rabbit anti-FITC (Invitrogen A889), and Mouse anti-Biotin (Invitrogen 03–3700) were used (1:400). Fluorescently conjugated secondaries, Alexa 555, 488, and 647, from ThermoFisher were used (1:400). See Table S1 for a complete list of reagents used (Table S1). Mouse anti-dorsal 7A4 (DSHB, (Whalen and Steward 1993)) was used for DL antibody staining (1:10), following the same protocol as FISH. Embryos that were the right stage based on visualization of a nuclear stain (DAPI) were imaged and staging was confirmed by *zen* and *ind* expression.

### Western Blot

For western blot analysis, embryos were collected and staged under white light. Embryos at mid nc14 were then added to a tube containing SDS buffer, ruptured using a fine needle, and homogenized. Embryo extracts were then run on a discontinuous SDS-PAGE gel and transferred to 0.45um Immobilon-P PVDF. Chemiluminescent detection was performed using a DL antibody (anti-Dorsal 7A4, DSHB, (Whalen and Steward 1993)). Beta-tubulin (E7, DSHB, (Chu and Klymkowsky 1989)) antibody was used as a loading control.

### Quantification and Statistical Analysis

To quantify the number of MS2 foci, or spots, in the images/movies captured, three custom MATLAB functions were used. The first function opens the image, including relevant metadata, and performs the spot detection. First “salt and pepper” noise is removed using a median filter. The background is subtracted by using a median filter over a larger area to blur the image, and then subtracting the blurred image from the original. After background subtraction, the image is blurred with a Gaussian filter. The image is then thresholded by a user defined threshold, tiny objects of only one pixel are removed, and objects detected on the edge are removed. The entire embryo is segmented by projecting all the time points together, blurring the image with a Gaussian filter, and using thresholding. The detected embryo is then morphologically closed to smooth the edges and small objects less than 100 pixels are discarded so only one object, the embryo, is detected. Any spot detected outside the embryo is removed. We observed that the background intensity of nuclei increased over time, and so to account for this, we increased the threshold by a small amount using a user defined value that increases logarithmically during nc14. To increase the detection of spots, we segmented the unprocessed image a second time using a user defined threshold and retained only the spots detected with both thresholds. The algorithm works by setting the first threshold low, detecting both real spots and noise, and then removing the noise based on a second, higher threshold. The centroid coordinates for these spots are saved for further analysis.

A second function displays the images in a GUI and allows overlapping the mask of the segmented foci on the image. In addition to overlaying, the centroids can be used to plot points at the detected spots. These were used to evaluate the success of the thresholding. Comparable imaging conditions used the same empirically determined threshold. Specifically, when comparing dark and light or *dl-LEXY* and *dl-BLID* the same threshold was used. The threshold was only changed for different MS2 signals (i.e. *sna* versus *sog*) or different lengths of imaging (i.e. nc12-nc14 versus 25 min of nc14). A third function was used to quantify the number of spots and plot the results. To plot the averages, the data was interpolated using MATLAB’s built in interp1 function and the Modified Akima cubic Hermite interpolation method. The mean number of spots and standard error of the mean were calculated from the interpolated data and plotted. In addition, this function also approximated the area of expression. This was done by concatenating all the centroids in given time windows corresponding to nc13 or the first 100 time points of nc14, and removing spots that were two median absolute deviations (MAD) from the median for the centroids of all spots detected. We used a conservative approach because this removed points that tended to be isolated, were not detected in multiple frames, or were actually noise and not a real spot. To determine the area, a convex hull was drawn around the remaining points and the area for this convex hull was determined. The area at nc13 was then subtracted from the area at nc14 to determine the change in nc13 to nc14 and account for potential discrepancies in the orientation of the embryo. This was only done for *sog-MS2* and *zen-MS2*.

To determine if the differences in area were statistically significant, we performed one way ANOVA and Tukey’s honestly significant difference (HSD) test for multiple comparisons to compare *dl-LEXY* dark, *dl-LEXY* light, *dl-BLID* dark, and *dl-BLID* light. We performed this analysis for the areas determined for both *sog-MS2* and *zen-MS2*. A p-value less than 0.05 was considered statistically significant.

To quantify the DL gradient in fixed samples, images were inputted into a previous developed function which measures the intensity of DL along the DV axis and fits the DL gradient to a gaussian function (Trisnadi et al. 2013).

To determine the sample size, a power analysis was done for each experiment. For the number of MS2 spots in *sna-MS2* a power analysis using a predicted mean of 200 spots, standard deviation of 40, power of 0.80, and significance value of 0.05 for a two sample t-test was done in MATLAB using sampsizepwr. This resulted in a sample size of n = 3. For the change in area of *sog-MS2* and *zen-MS2*, this procedure was repeated using a mean area of 2000, standard deviation of 1500, power of 0.80, and significance value of 0.05 for a two sample t-test. This also resulted in a sample size of n = 3. Similarly, we performed this analysis for the quantification of the DL gradient using a mean amplitude of 100, standard deviation of 25, power of 0.80, and significance value of 0.05 for a two sample t-test. This resulted in a sample size of n = 6.

## ACKNOWLEDGMENTS

We are grateful to Hernan Garcia and Mike Levine for sharing fly stocks and plasmids, and to Vince Stepanik and Lelise Dunipace for technical support and comments on the manuscript. We also thank Virginia Pimmett, Antonio Trullo, and Mounia Lagha for helpful discussions.

## DATA AVAILABILITY

*Drosophila* strains and other reagents generated in this study will be available upon request from the lead contact, or the commercial sources listed in Table S1.

A github repository with the codes for quantitative analyses were generated and are publicly available: (https://github.com/StathopoulosLab/MS2_quantification). Any additional information required to reanalyze the data shown in this paper is available from the lead contact upon request.

## AUTHOR CONTRIBUTIONS

A.S. and J.M. conceived the project. J.M. planned the experimental approach. A.S. directed the project. J.M. performed all experiments including the computational work. J.M. analyzed the data with input from A.S. The manuscript was written by J.M. and A.S.

## SUPPLEMENTAL INFORMATION TITLES AND LEGENDS

### Supplemental Figures and Legends

**Figure S1**. MS2 reporter constructs. Related to Figure 2 and 3.

**Figure S2**. Variability in starting number of transcription sites for *sna-MS2*. Related to Figure 2.

**Figure S3**. In both *dl-BLID* and *dl-LEXY*, no change in the *zen-MS2* boundary is observed when comparing nc13 to nc14. Related to Figure 3.

**Figure S4**. Twi is not exported with DL-LEXY, suggesting cofactors are not sequestered in the cytoplasm.

**Figure S5**. Peak levels of DL are lower in *dl NES S>A* than in Control embryos. Related to Figure 6.

### Supplemental Table

**Table S1**. A list of the *Drosophila melanogaster* lines, primers, plasmids, reagents, and software used in this study.

### Movie legends

**Movie 1.** Blue light-induced export of DL-mCherry-LEXY is rapid and reversible. Related to Figure 1. Blue bar in the upper right corner represents frames under blue light in this and all subsequent movies.

**Movie 2.** *sna-MS2* in *dl-LEXY* and *dl-BLID* in the dark, with 10 min of blue light, and with 20 min of blue light. Related to Figure 2. Nascent transcription was only detected above a certain threshold and was marked by circles in this and all following movies.

**Movie 3.** *sog-MS2* in *dl-LEXY* and *dl-BLID* in the dark and with blue light throughout nc13 and 14. Related to Figure 3.

**Movie 4.** DL-mCh-LEXY and DL-mCh-BLID in the dark, with 10 min of blue light and with 20 min of blue light. Related to Figure 4.

**Movie 5.** FRAP of DL-mCh-LEXY in an ROI under blue light. Related to Figure 5.

**Movie 6.** *zen-MS2* in *dl-LEXY* and *dl-BLID* in the dark and with blue light throughout nc13 and 14. Related to Supplementary Figure 3.

**Movie 7.** Twi-LlamaTag bound to mCherry with 10 min of blue light. Related to Supplementary Figure 4.

**Figure S1.**
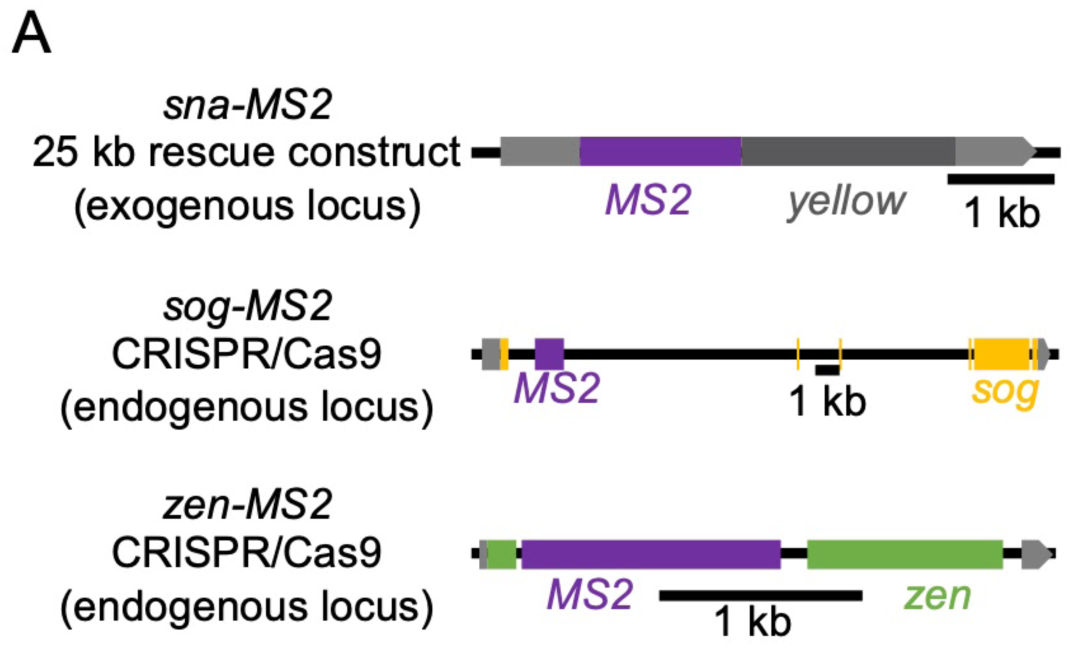
MS2 reporter constructs. (A) The MS2 constructs used for observing active transcription live. The *sna-MS2* line was from a previously published study (Bothma et al. 2015) whereas the *sog-MS2* and *zen-MS2* reporters were created in this study. The *sna-MS2* reporter is a large reporter construct (∼25 kB) inserted as an exogenous copy on the 3rd chromosome. Shown here is just the MS2 and exons of the *sna-MS2* reporter. The *sog-MS2* and *zen-MS2* are reporters with MS2 inserted at the endogenous loci using Crispr/Cas9 (see Methods).

**Figure S2.**
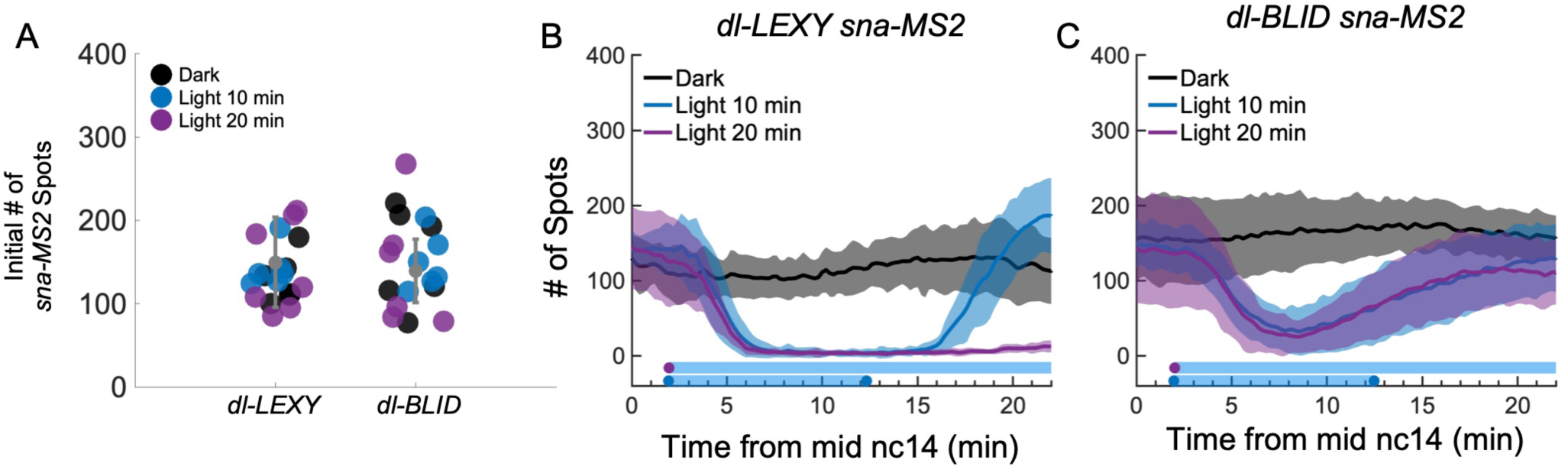
Variability in starting number of transcription sites for *sna-MS2*. (A) The starting number of spots for *dl-LEXY* and *dl-BLID* with spots color coded to match each condition: dark (black), light 10 min (blue), and light 20 min (purple). The number of spots did not have a statistically significant difference between the means (p = 0.44, Tukey’s HSD for multiple comparisons after performing one way ANOVA, in black mean ± s.d., for *dl-LEXY* n = 19 and for *dl-BLID* n = 18.). **(B,C)** The unnormalized mean number of spots for *dl-LEXY* (B) and *dl-BLID* (C) in the dark (black), with 10 min of blue light (blue), or 20 min of blue light (purple). The number of initial spots of *sna* active transcription can vary for multiple reasons. One reason is that both BLID and LEXY are leaky in the dark, and this could lead to differences in DL levels which affect the number of active sites. In addition, since DL-LEXY and DL-BLID have different effects on *sna* under blue light, it is likely that this leaky behavior would result in differences in *sna*. These embryos are staged by tracking the cellularization front and imaging begins when the front is halfway down the length of the nuclei (mid to late nc14). This staging technique is imperfect and also leads to variability in starting number of spots for *sna*, as *sna* sites of active transcription are known to vary over time (Bothma et al. 2015). Embryos were collected from the same respective cages either on the same day or subsequent days.

**Figure S3.**
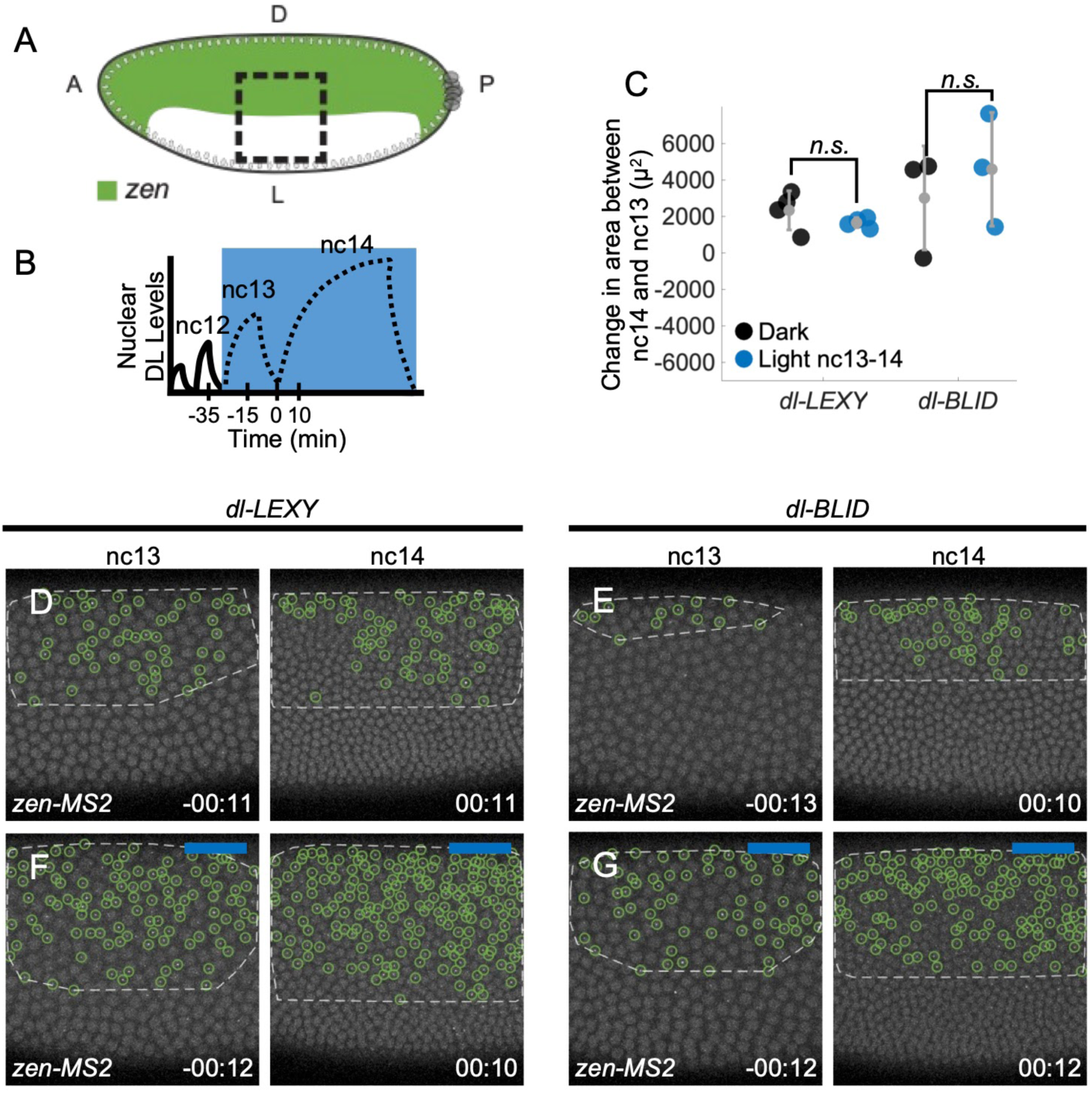
In both *dl-BLID* and *dl-LEXY*, no change in the *zen-MS2* boundary is observed when comparing nc13 to nc14. (A) A schematic of *zen* expression with the field of view and blue light illumination window marked by the dashed black box. **(B)** The blue light illumination window from nc13 through the end of nc14. **(C)** Quantification of the change in area between nc14 and 13 for *zen-MS2* in *dl-LEXY* and *dl-BLID*. Black markers represent the dark and blue markers represent illumination from nc13-14 (in gray, mean ± s.d., n = 3 for *dl-BLID* and n = 4 for *dl-LEXY*). When comparing dark to light at nc13-14, the means are not significantly different, p = 0.96 for *dl-LEXY* light vs. dark and p = 0.77 for *dl-BLID* light vs. dark (Tukey’s HSD for multiple comparisons after performing one way ANOVA). **(D-G)** *zen-MS2* in *dl-LEXY* (D,F) and *dl-BLID* (E,G), when kept in the dark (D,F) or when illuminated continuously from nc13 through nc14 (F,G) at nc13 (–00:11, –00:12, and –00:13) and early-nc14 (00:10, 00:11, and 00:12). Variability in the change in area between nc14 and 13 is higher in *dl-BLID* and it appears that between nc13 and nc14 the *zen* domain is expanding. However, this expansion and the variability result in no statistically significant difference between the light and the dark. This may have to do with how *zen* senses DL. In *dl-LEXY* the low levels of DL are likely sufficient for repressing *zen*, but in *dl-BLID*, the leaky nature of *dl-BLID* might result in different levels of DL and *zen* might be sensitive to this. Embryos of a certain genotype were collected from the same cage either on the same day or subsequent days.

**Figure S4.**
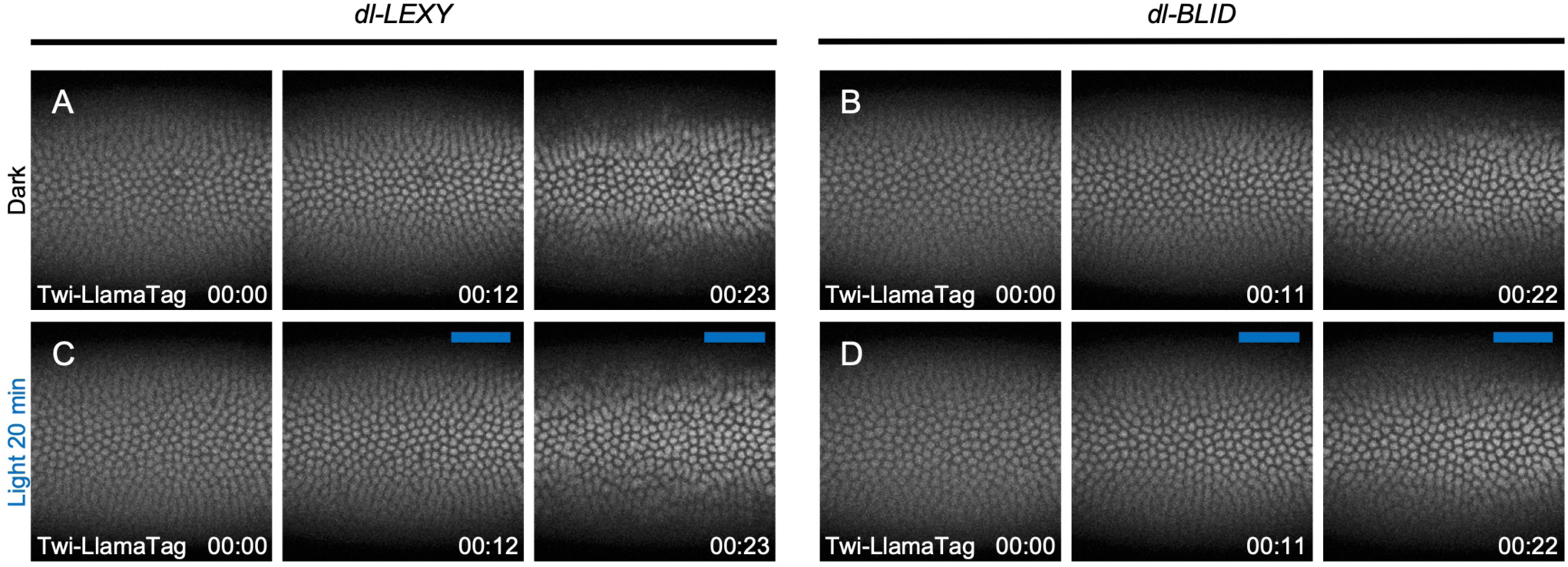
Twi levels do not change at late nc14 under blue light in *dl-LEXY* or *dl-BLID*. See also Movie 7. (A, B) Twi-Llama-Tag bound to mCherry in *dl-LEXY* (A) and *dl-BLID* (B) in the dark. **(C,D)** Twi-Llama-Tag bound to mCherry in *dl-LEXY* (C 00:12 and 00:23) and *dl-BLID* (D 00:11 and 00:22) after blue light exposure. In both *dl-LEXY* and *dl-BLID*, the levels of Twi do not change in blue light and levels are comparable between the two lines. Thus, a change in Twi levels does not explain the difference in *sna* expression bewteen *dl-LEXY* and *dl-BLID* under blue light. In addition, since little to no effect on nuclear levels of Twi was observed when embryos were illuminated with light and Twi levels did not increase in the cytoplasm, this result suggests that export of DL from the nucleus does not drag out other transcriptional cofactors to inhibit *sna* expression indirectly. For each condition, n = 3, and embryos were collected from the same cage either on the same day or subsequent days.

**Figure S5.**
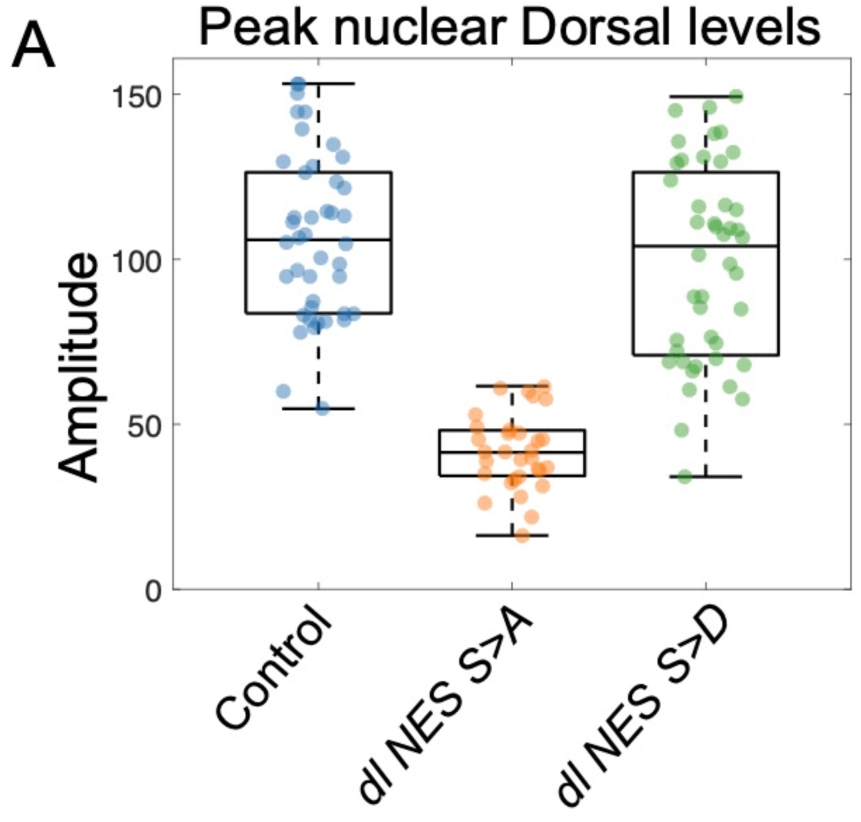
Peak levels of DL are lower in *dl NES S>A* than in Control embryos. (A) A plot of the amplitudes or peak levels of DL for the Control (*dl-Venus; blue*), *dl NES S>A (orange)*, and *dl NES S>D (green)*. In black is a box-and-whiskers plot showing (from bottom to top) the minimum, first quartile, median, third quartile, and the maximum. The amplitude was determined by fitting the nuclear levels of the DL gradient to a Gaussian function. The amplitude reflects the peak levels of DL, or the levels of the ventral most nucleus, where DL levels are highest. The levels in *dl NES S>A* are lower than the Control or *dl NES S>D* (p = 8.6 * 10^-20^ and 7.8 * 10^-17^, Tukey’s HSD for multiple comparisons after performing one way ANOVA). For Control n = 42, *dl NES S>A* n = 31, and for *dl NES S>D* n = 44. Embryos of a certain genotype were collected from the same cage either on the same day or subsequent days.

