## Supplemental information for "Target gene responses differ when transcription factor levels are acutely decreased by nuclear export versus degradation"

**Figure S1** related to Figure 1

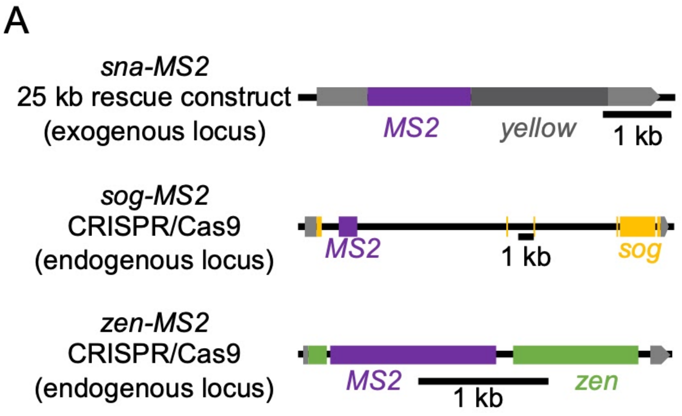


**Figure S1. MS2 reporter constructs. (A)** The MS2 constructs used for observing active transcription live. The *sna-MS2* line was from a previously published study (Bothma et al. 2015) whereas the *sog-MS2* and *zen-MS2* reporters were created in this study. The *sna-MS2* reporter is a large reporter construct (~25 kB) inserted as an exogenous copy on the 3rd chromosome. Shown here is just the MS2 and exons of the *sna-MS2* reporter. The *sog-MS2* and *zen-MS2* are reporters with MS2 inserted at the endogenous loci using Crispr/Cas9 (see Methods).

**Figure S2** related to Figures 1, 2, and 3

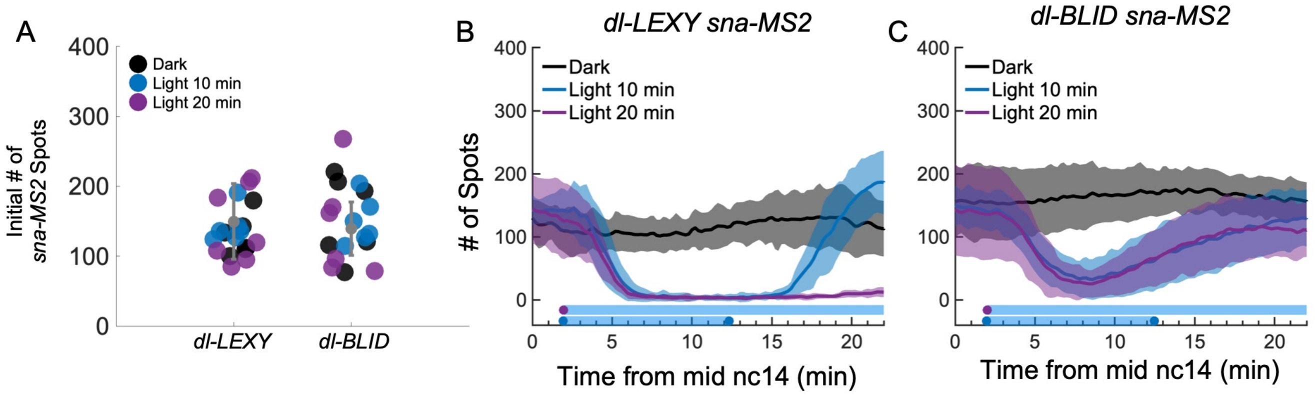


**Figure S2. Variability in starting number of transcription sites for *sna-MS2*. (A)** The starting number of spots for *dl-LEXY* and *dl-BLID* with spots color coded to match each condition: dark (black), light 10 min (blue), and light 20 min (purple). The number of spots did not have a statistically significant difference between the means ($p = 0.44$, Tukey's HSD for multiple comparisons after performing one way ANOVA, in black mean ± s.d., for *dl-LEXY* n = 19 and for *dl-BLID* n = 18.). **(B,C)** The unnormalized mean number of spots for *dl-LEXY* (B) and *dl-BLID* (C) in the dark (black), with 10 min of blue light (blue), or 20 min of blue light (purple). The number of initial spots of *sna* active transcription can vary for multiple reasons. One reason is that both BLID and LEXY are leaky in the dark, and this could lead to differences in DL levels which affect the number of active sites. In addition, since DL-LEXY and DL-BLID have different effects on *sna* under blue light, it is likely that this leaky behavior would result in differences in *sna*. These embryos are staged by tracking the cellularization front and imaging begins when the front is halfway down the length of the nuclei (mid to late nc14). This staging technique is imperfect and also leads to variability in starting number of spots for *sna*, as *sna* sites of active transcription are known to vary over time (Bothma et al. 2015). Embryos were collected from the same respective cages either on the same day or subsequent days.

### Figure S3

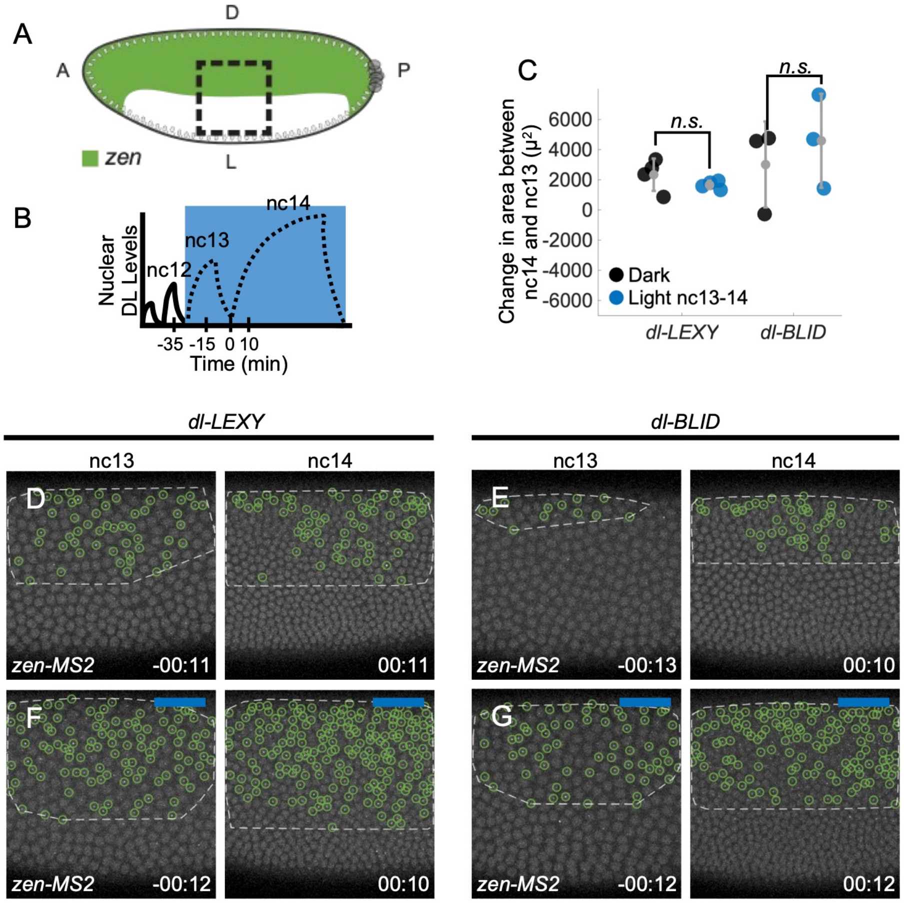


**Figure S3. In both *dl-BLID* and *dl-LEXY*, no change in the *zen-MS2* boundary is observed when comparing nc13 to nc14. (A)** A schematic of *zen* expression with the field of view and blue light illumination window marked by the dashed black box. **(B)** The blue light illumination window from nc13 through the end of nc14. **(C)** Quantification of the change in area between nc14 and 13 for *zen-MS2* in *dl-LEXY* and *dl-BLID*. Black markers represent the dark and blue markers represent illumination from nc13-14 (in gray, mean ± s.d., n = 3 for *dl-BLID* and n = 4 for *dl-LEXY*). When comparing dark to light at nc13-14, the means are not significantly different, p = 0.96 for *dl-LEXY* light vs. dark and p = 0.77 for *dl-BLID* light vs. dark (Tukey's HSD for multiple comparisons after performing one way ANOVA). **(D-G)** *zen-MS2* in *dl-LEXY* (D,F) and *dl-BLID* (E,G), when kept in the dark (D,F) or when illuminated continuously from nc13 through nc14 (F,G) at nc13 (-00:11, -00:12, and -00:13) and early-nc14 (00:10, 00:11, and 00:12). Variability in the change in area between nc14 and 13 is higher in *dl-BLID* and it appears that between nc13 and nc14 the *zen* domain is expanding. However, this expansion and the variability result in no statistically significant difference between the light and the dark. This may have to do with how *zen* senses DL. In *dl-LEXY* the low levels of DL are likely sufficient for repressing *zen*, but in *dl-BLID*, the leaky nature of *dl-BLID* might result in different levels of DL and *zen* might be sensitive to this. Embryos of a certain genotype were collected from the same cage either on the same day or subsequent days.

#### Figure S4

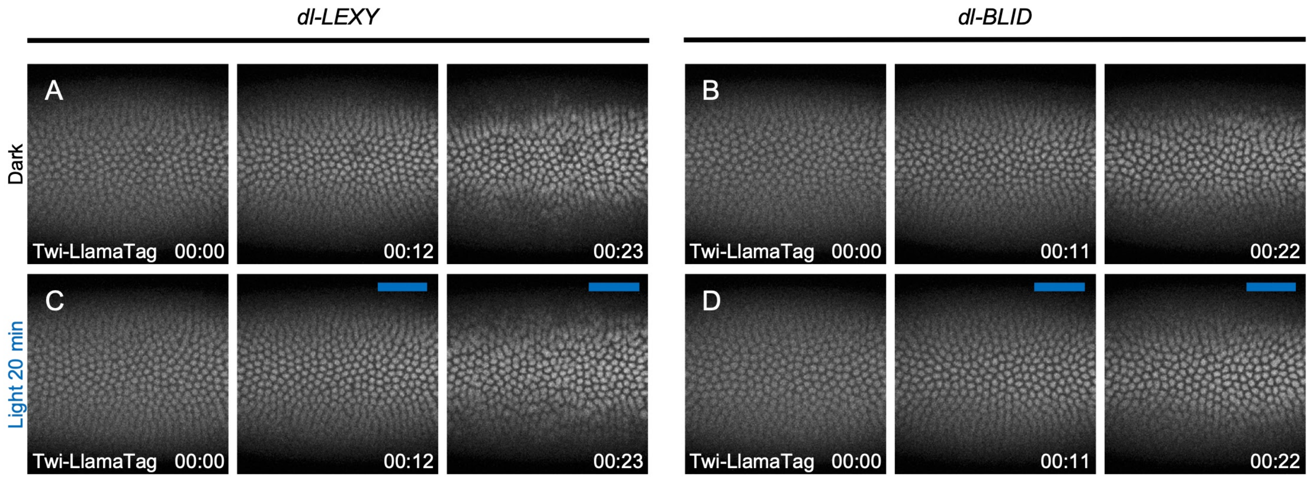


**Figure S4. Twi levels do not change at late nc14 under blue light in *dl-LEXY* or *dl-BLID*. See also Movie 7. (A, B)** Twi-Llama-Tag bound to mCherry in *dl-LEXY* (A) and *dl-BLID* (B) in the dark. **(C,D)** Twi-Llama-Tag bound to mCherry in *dl-LEXY* (C 00:12 and 00:23) and *dl-BLID* (D 00:11 and 00:22) after blue light exposure. In both *dl-LEXY* and *dl-BLID*, the levels of Twi do not change in blue light and levels are comparable between the two lines. Thus, a change in Twi levels does not explain the difference in *sna* expression bewteen *dl-LEXY* and *dl-BLID* under blue light. In addition, since little to no effect on nuclear levels of Twi was observed when embryos were illuminated with light and Twi levels did not increase in the cytoplasm, this result suggests that export of DL from the nucleus does not drag out other transcriptional cofactors to inhibit *sna* expression indirectly. For each condition, n = 3, and embryos were collected from the same cage either on the same day or subsequent days.

#### Figure S5

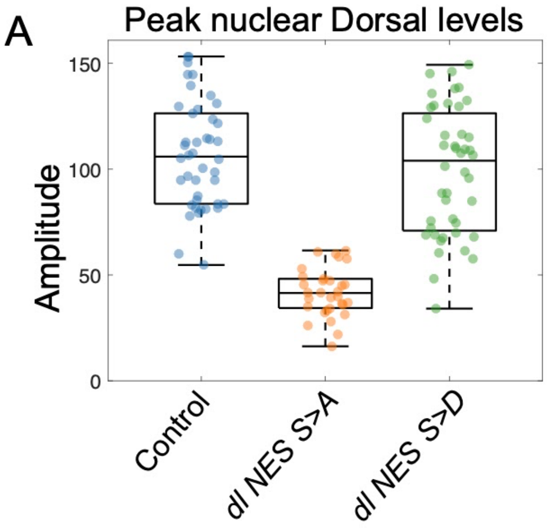


**Figure S5. Peak levels of DL are lower in *dl NES S>A* than in Control embryos. (A)** A plot of the amplitudes or peak levels of DL for the Control (*dl-Venus; blue*), *dl NES S>A (orange)*, and *dl NES S>D (green)*. In black is a box-and-whiskers plot showing (from bottom to top) the minimum, first quartile, median, third quartile, and the maximum. The amplitude was determined by fitting the nuclear levels of the DL gradient to a Gaussian function. The amplitude reflects the peak levels of DL, or the levels of the ventral most nucleus, where DL levels are highest. The levels in *dl NES S>A* are lower than the Control or *dl NES S>D* ($p = 8.6 * 10^{-20}$ and $7.8 * 10^{-17}$, Tukey's HSD for multiple comparisons after performing one way ANOVA). For Control n = 42, *dl NES S>A* n = 31, and for *dl NES S>D* n = 44. Embryos of a certain genotype were collected from the same cage either on the same day or subsequent days.
