## Supplementary material for "Target gene responses differ when transcription factor levels are acutely decreased by nuclear export versus degradation": Table S1

| REAGENT or RESOURCE | SOURCE | IDENTIFIER |
|---|---|---|
| **Antibodies** | | |
| Mouse Anti-Dorsal | Developmental Studies Hybridoma Bank (DSHB, (Whalen and Steward 1993) | Cat# anti-dorsal 7A4; RRID: AB_528204 |
| Mouse Anti-Beta tubulin (E7) | Developmental Studies Hybridoma Bank (DSHB, (Chu and Klymkowsky 1989) | Cat# E7; RRID: AB_528499 |
| Sheep Anti-Digoxigenin Polyclonal Antibody | Thermo Fisher Scientific | Cat# PA1-85378; RRID:AB_930545 |
| Rabbit Anti Fluorescein isothiocyanate | Thermo Fisher Scientific | Cat# A889; RRID: AB_221561 |
| Mouse Anti Biotin | Thermo Fisher Scientific | Cat# 03-3700; RRID: AB_2532265 |
| Alexa Fluor 488 donkey anti-mouse | Thermo Fisher Scientific | Cat# A21202; RRID: AB_141607 |
| Alexa Fluor 488 donkey anti-sheep | Thermo Fisher Scientific | Cat# A11015; RRID: AB_2534082 |
| Alexa Fluor 555 donkey anti-mouse | Thermo Fisher Scientific | Cat# A31570; RRID: AB_2536180 |
| Alexa Fluor 647 donkey anti-rabbit | Thermo Fisher Scientific | Cat# A31573; RRID: AB_2536183 |

| | | |
|---|---|---|
| **Chemicals, Peptides, and Recombinant Proteins** | | |
| Digoxigenin labeled nucleotides | Roche | Cat# 11277073910 |
| Biotin labeled nucleotides | Roche | Cat# 11685597910 |
| Fluorescein isothiocyanate labeled nucleotides | Roche | Cat# 11685619910 |
| **Experimental Models: Organisms/Strains** | | |
| *D. melanogaster*: *w; dl-LEXY/CyO; PrDr/TM3* | This study | N/A |
| *D. melanogaster*: *w; dl-BLID/CyO; PrDr/TM3* | Irizarry et al.(Irizarry et al. 2020) | N/A |
| *D. melanogaster*: *Sp/Cyo; MCP-mCherry (w+, NLS)/TM3* | Bothma et al.(Bothma et al. 2018) | N/A |
| *D. melanogaster*: *w; dl-LEXY/CyO; MCP-mCherry (w+, NLS)/TM3* | This study | N/A |
| *D. melanogaster*: *w; dl-BLID/CyO; MCP-mCherry (w+, NLS)/TM3* | This study | N/A |
| *D. melanogaster*: w; *dl-mCherry-LEXY/CyO* | This study | N/A |
| *D. melanogaster*: w; *dl-mCherry-BLID/CyO* | Irizarry et al.(Irizarry et al. 2020) | N/A |
| *D. melanogaster*: *sna-MS2 BAC (III)* | Bothma et al.(Bothma et al. 2015) | N/A |

| | | |
|---|---|---|
| *D. melanogaster*: *sog-MS2 (I); Sp/CyO* | This study | N/A |
| *D. melanogaster*: *Sp/CyO; zen-MS2/TM3 (III)* | This study | N/A |
| *D. melanogaster*: *y2 cho2 v1 P{nos-phiC31\int.NLS}X; P{dl-gRNA}attP2 (III)* | Irizarry et al.(Irizarry et al. 2020) | N/A |
| *D. melanogaster*: *y2 cho2 v1; Sp/CyO, P{nos-Cas9, y+, v+}2A* | NIG-FLY | Cat#CAS-0004 |
| *D. melanogaster*: *dl-Venus (III)* | Reeves et al.(Reeves et al. 2012) | N/A |
| *D. melanogaster*: *$dl^1/dl^4$; dl-Venus (III)* | Reeves et al.(Reeves et al. 2012) | N/A |
| **Oligonucleotides** | | |
| Primer: Dorsal Fusion Forward CACAAAACATACCGCCCATT | This study | N/A |
| Primer: Dorsal Fusion Reverse CGCTTCCTCCCGTGGATATGGACAGGTTCG | This study | N/A |
| Primer: LEXY Fusion Forward CATATCCACGGGAGGAAGCGGAGGAAGCGGA | This study | N/A |
| Primer: LEXY Fusion Reverse GAAAAAGGTATCAATCCAGGTTCAGGTCGGCC | This study | N/A |
| Primer: 3'UTR Fusion Forward CCTGGATTGATACCTTTTTCACAACGAACCAG | This study | N/A |

| | | |
|---|---|---|
| Primer: 3'UTR Fusion Reverse AAGTGGGTGGGCAGCTTATC | This study | N/A |
| Primer: mCherry LEXY Fusion Reverse CGCTTCCTCCCTTGTACAGCTCGTCCATGC | This study | N/A |
| Primer: mCherry LEXY Fusion Forward GCTGTACAAGGGAGGAAGCGGAGGAAGCGGA | This study | N/A |
| Primer: sog left homology arm fragment 1 Forward TTTGGGTTCGGCATTTAGAG | This study | N/A |
| Primer: sog left homology arm fragment 1 Reverse ACAGTTAATATCCCTTTAACTAAAGATTTTGAC | This study | N/A |
| Primer: sog left homology arm fragment 2 Forward AGTTAAAGGGATATTAACTGTTGCCTGTTGC | This study | N/A |
| Primer: sog left homology arm fragment 2 Reverse CGCCCGTTTCTGCATTATGA | This study | N/A |
| Primer: sog left homology arm full fragment Forward ATCGCTAGCCGTTGTCGTAATCCCTCCTC | This study | N/A |
| Primer: sog left homology arm full fragment Reverse ATGATTAATCCTAGGACTGCGGCCGCCGTATCAACTAAGCCCTAAC | This study | N/A |
| Primer: sog right homology arm Forward ATTGGCGCGCCACGTAGATCCCGGGATTTGTG | This study | N/A |

| | | |
|---|---|---|
| Primer: sog right homology arm Reverse ATTCTCGAGACATCCACCATCCACCACAT | This study | N/A |
| Primer: zen gRNA1 pCFD5 Forward GCGGCCCGGGTTCGATTCCCGGCCGATGCATTGTAGGTAGACGACATAACGTTTTAGAGCTAGAAATAGCAAG | This study | N/A |
| Primer: zen gRNA2 pCFD5 Reverse ATTTTAACTTGCTATTTCTAGCTCTAAAACATGACACATGCTGCTGATGCTGCACCAGCCGGGAATCGAACCC | This study | N/A |
| Primer: zen left homology arm fragment 1 Forward CATACTCACACATGCCTGCC | This study | N/A |
| Primer: zen left homology arm fragment 1 Reverse GACATAACAGATCAATTCCGCCGACGCAT | This study | N/A |
| Primer: zen left homology arm fragment 2 Forward GCGGAATTGATCTGTTATGTCGTCTACCTACAAG | This study | N/A |
| Primer: zen left homology arm fragment 2 Reverse TTAAGCTTCACCCTCTGCGA | This study | N/A |
| Primer: zen left homology arm full fragment Forward ATCGCTAGCCAATTGTGCACAGTGACCCA | This study | N/A |
| Primer: zen left homology arm full fragment Reverse ATGCATATGCCTAGGACTGCGGCCGCTGCTGATATTCGATTTTTGATCT | This study | N/A |
| Primer: zen right homology arm fragment 1 Forward CAGCGACGAAGGGATTTACG | This study | N/A |

| | | |
|---|---|---|
| Primer: zen right homology arm fragment 1 Reverse AGATTCTCGTTCGATGACACATGCTGCTGAT | This study | N/A |
| Primer: zen right homology arm fragment 2 Forward TGTGTCATCGAACGAGAATCTGCCATCTCAG | This study | N/A |
| Primer: zen right homology arm fragment 2 Reverse CAGCAAGCAACAAGGAGTCA | This study | N/A |
| Primer: zen right homology arm full fragment Forward ATTGGCGCGCCATAAACTAATTAGTTGTTTACCCGC | This study | N/A |
| Primer: zen right homology arm full fragment Reverse ATTCTCGAGAGACGGTTGCTTAGCTCCAA | This study | N/A |
| **Recombinant DNA** | | |
| pHD-DsRed | Gratz et al.(Gratz et al. 2014) | Addgene 51434 |
| Plasmid: LEXY (codon optimized for *D. melanogaster)* | This study | N/A |
| Plasmid: dl-LEXY, DsRed homologous repair | This study | N/A |
| Plasmid: dl-mCh-LEXY, DsRed homologous repair | This study | N/A |
| Plasmid: sog-MS2 gRNA | Dunipace et al.(Dunipace, Ákos, and Stathopoulos 2019) | N/A |

| | | |
|---|---|---|
| Plasmid: sog-MS2, DsRed homologous repair | This study | N/A |
| Plasmid: zen-MS2 gRNA | This study | N/A |
| Plasmid: zen-MS2, DsRed homologous repair | This study | N/A |
| **Software and Algorithms** | | |
| Zen 3.0 (Blue edition) | Zeiss | N/A |
| Fiji/ImageJ | Schindelin et al.(Schindelin et al. 2012) | https://imagej.nih.gov/ij/ |
| MS2_quantification | This study | https://github.com/StathopoulosLab/MS2_quantification |
| run_analyze_xs | Trisnadi et al. (Trisnadi et al. 2013) | https://www.sciencedirect.com/science/article/pii/S1046202312002629?via%3Dihub |
